# Default mode network shows distinct emotional and contextual responses yet common effects of retrieval demands across tasks

**DOI:** 10.1101/2023.09.30.560279

**Authors:** Nicholas E. Souter, Antonia de Freitas, Meichao Zhang, Ximing Shao, Tirso Rene del Jesus Gonzalez Alam, Haakon Engen, Jonathan Smallwood, Katya Krieger-Redwood, Elizabeth Jefferies

## Abstract

The default mode network (DMN) lies towards the heteromodal end of the principal gradient of intrinsic connectivity, maximally separated from sensory-motor cortex. It supports memory-based cognition, including the capacity to retrieve conceptual and evaluative information from sensory inputs, and to generate meaningful states internally; however, the functional organisation of DMN that can support these distinct modes of retrieval remains unclear. We used fMRI to examine whether activation within subsystems of DMN differed as a function of retrieval demands, or the type of information to be retrieved, or both. In a picture association task, participants retrieved two types of semantic features about contexts and emotions: in the generate phase, these associations were retrieved from a novel picture, while in a switch phase, participants retrieved a new association for the same image. Semantic context and emotion trials were associated with dissociable DMN subnetworks, indicating that a key dimension of DMN organisation relates to the type of information being accessed. The fronto-temporal and medial temporal DMN showed a preference for emotional and contextual associations, respectively. Relative to the generate phase, the switch phase recruited clusters closer to the heteromodal apex of the principal gradient – a cortical hierarchy separating unimodal and heteromodal regions. There were no differences in this effect between association types. Instead, memory switching was associated with a distinct subnetwork associated with controlled internal cognition. These findings delineate *distinct* patterns of DMN recruitment for different kinds of associations yet *common* responses across tasks that reflect retrieval demands.

**Key points:**

1. Retrieval of contextual and emotional features relies on distinct default mode subnetworks.
2. Novelty of visual input has an equivalent effect across default mode subnetworks.
3. Secondary associations retrieved from re-presented pictures activate heteromodal end of the principal gradient.

## 1. Introduction

The default mode network (DMN) is a large-scale distributed network which frequently shows task-related deactivation (Raichle, 2015) yet is also associated with aspects of cognition that are dependent on memory (Murphy et al., 2018; Zhang et al., 2022). It is thought to support diverse tasks relating to social cognition, episodic recall, semantic retrieval, and emotion induction (Mancuso et al., 2022). In such domains, DMN is thought to support our interpretation of external events and the distillation of diverse features (Lanzoni et al., 2020), as well as the ability to generate cognitive states that are decoupled from the external world (Smallwood et al., 2021). However, the functional organisation of this network remains unclear, since its activation may be modulated according to the type of information being retrieved, and/or the retrieval demands associated with a particular task.

DMN regions are thought to be maximally distant from sensory-motor cortex on a cortical hierarchy. This topographical organisation is captured by the principal gradient of intrinsic connectivity, which is the dimension of whole-brain connectivity that explains the most variance (Margulies et al., 2016). The principal gradient reveals maximal separation of connectivity patterns between unimodal and heteromodal regions and can also explain the order of large-scale networks along the cortical surface, from sensory-motor regions, through attention networks and the frontoparietal control network, to DMN. This separation is thought to allow DMN to support both perceptually-decoupled and abstract thought, since both involve informational states that are at odds with the changing environment (Gordon et al., 2020; Murphy et al., 2018; Smallwood et al., 2021). For example, from picture cues, we may access information about abstract categories that allow us to evaluate and make sense of our experiences and retrieve past events that are no longer taking place (using sensory to DMN pathways). We may also generate sensory-motor features relating to these concepts or past events, even when they do not overlap with features present in the external world (using DMN to sensory pathways).

The role DMN plays in these processes may be better understood by investigating how dissociable subnetworks support distinct types of memory-guided cognition. Andrews-Hanna et al. (2014) reported that in addition to a ‘core’ DMN network focussed on anterior and posterior cingulate cortex and angular gyrus, patterns of intrinsic connectivity reveal a medial temporal (MT) subnetwork (including retrosplenial cortex) and a lateral fronto-temporal (FT) subnetwork^1^ (including dorsomedial prefrontal cortex). The MT subnetwork includes aspects of the hippocampus and parahippocampal gyrus, implicated in mental scene construction (Sheldon & Levine, 2016). The FT subnetwork shows greater connectivity with the anterior temporal lobes (ATL; Andrews-Hanna et al., 2014), a brain region thought to provide a heteromodal semantic ‘hub’ (Chiou & Lambon Ralph, 2019; Lambon Ralph et al., 2017) which is also associated with processing valence (Juran et al., 2016; Spiers et al., 2017; Wang et al., 2019). These subnetworks are consequently associated with different aspects of memory: the MT network is associated with episodic recollections of specific experiences that are typically visuo-spatial in nature, while the FT network is implicated in semantic and social cognition, based on knowledge extracted across many experiences which is typically more abstract in nature (Andrews-Hanna and Grilli, 2021; Andrews-Hanna et al., 2014; Chiou et al., 2020; Gurguryan & Sheldon, 2019; Sheldon et al., 2019). Core DMN regions sit at points where the MT and FT subsystems are spatially interdigitated (Braga & Buckner, 2017; Yeo et al., 2011) and might help to draw together spatial, semantic, and valence information to form coherent patterns of cognition (Lanzoni et al., 2020).

Despite progress in characterising the functional organisation of DMN, important issues remain unresolved. The first concerns the specific task dimensions that separate MT and FT subsystems. While semantic cognition is often considered to involve abstract categories (e.g., types of emotions), we also have general visuo-spatial knowledge about typical scenes and events, acquired over our lifetime. Contrasts of semantic tasks probing contextual information about generic spatial-temporal associations versus valenced associations can establish whether MT and FT subnetworks support these distinct types of knowledge (scene construction versus evaluative categories). A second unresolved issue concerns how retrieval demands intersect with DMN subnetworks. Irrespective of the type of information being retrieved (e.g., contextual or emotional associations), retrieval tasks can vary in the cognitive resources that they utilise; for example, they might differ in their reliance on immediately available sensory-motor features versus representations from memory. Research to date has shown similar DMN recruitment for tasks based on meaning as opposed to individual perceptual features and for memory-guided 1-back decisions over 0-back trials (Murphy et al., 2018). Key DMN regions, including posterior cingulate and medial prefrontal cortex, are also implicated in the disambiguation of degraded ambiguous images when they are re-presented and recognised (González-García et al., 2018), suggesting the heteromodal end of the principal gradient supports the processing of more familiar images that have become meaningful. Nevertheless, very little research has investigated whether these kinds of retrieval demands equivalently modulate the response of distinct DMN subsystems. For example, FT might be more coupled to visual cortex (to allow semantic access from vision for novel images) while MT might be more perceptually-decoupled and relevant to patterns of retrieval that rely largely on internal processes (Chiou et al., 2020; Zhang et al., 2022). Alternatively, manipulations of perceptual novelty or the task-relevance of visual input might alter DMN recruitment in a way that is orthogonal to the MT/FT distinction. Internal mind-wandering relies on conceptual and evaluative information (Faber & D’Mello, 2018; Smallwood et al., 2016), suggesting that the FT subsystem might be implicated in both perceptually-generated and decoupled states.

Further debate concerns the relationship between DMN and cognitive control networks. DMN regions can support memory-based cognition when executive demands are high (Brown et al., 2019; Murphy et al., 2018; 2019) and DMN is implicated in goal maintenance during controlled semantic retrieval (Wang et al., 2021). This functional similarity between DMN and heteromodal control networks is captured by the principal gradient (Margulies et al., 2016). Frontotemporal control regions also form alliances that reflect task demands (Cole et al., 2013; Gonzalez Alam et al., 2021; Niendam et al., 2012; Spreng et al., 2013); they are recruited with attention networks to form the multiple demand network (MDN; Duncan, 2001; 2010; Fedorenko et al., 2013; Hugdahl et al., 2015), while a semantic control network (SCN) supports the conceptual retrieval of non-dominant aspects of knowledge relevant to the current task (Noonan et al., 2013; Jackson, 2021). SCN is situated at the intersection of frontoparietal control regions and DMN (Chiou et al., 2022; Davey et al., 2016; Wang et al., 2020), and is functionally and spatially distinct from MDN (Gao et al., 2021; Humphreys et al., 2015). This dissociation somewhat resembles parcellations of control networks based on intrinsic connectivity: the ‘control A’ subnetwork^2^ couples with the externally-oriented dorsal attention network (DAN), while ‘control B’ couples with DMN (Dixon et al., 2018; Yin et al., 2022). However, little is known about how distinctions within control networks relate to MT and FT DMN subdivisions (see Vatansever et al., 2021).

Here, we used functional magnetic resonance imaging (fMRI) to characterise the contribution of DMN subsystems and control networks to the retrieval of emotional associations and meaning-based contexts from picture cues. While both tasks were reliant on semantic information, they may preferentially activate FT and MT subnetworks, respectively, if these subsystems are sensitive to the *type* of information being retrieved. Associations were made twice for each picture, allowing us to manipulate the relevance of visual input and memory-based processes to retrieval (via the novelty of visual input, following González-García et al., 2018). The ‘*generate’* phase required participants to identify an association from a novel picture, while the ‘*switch’* phase required a *new* association from the same picture. We situated responses associated with each phase on the principal gradient, to test whether the switch phase elicited stronger recruitment of heteromodal cortex. We also investigated whether responses in FT and MT DMN were modulated by the differing demands of *generate* versus *switch* phases. Alternatively, this manipulation may be largely independent of association type and instead reflected in control-relevant DMN regions (e.g., control B).

## 2. Materials & Methods

### 2.1. Participants

Participants were right-handed, aged 18 to 35, with normal or corrected to normal vision, no history of neurological disorder, and no current psychiatric disorder. Participants were students at the University of York, recruited through word of mouth and participant pools, and paid for their time or awarded course credit. One participant was excluded as they reported no association was retrieved on 52% of trials (see Section 2.4). The final sample consisted of 32 participants (24 female) with a mean age of 20.1 (SD = 2.4). Ethical approval was granted by the York Neuroimaging Centre (date: 23/06/2021, project ID: P1446). Informed consent was obtained from all participants prior to participation.

### 2.2. Design

A within-subjects design was used. Each participant retrieved *emotion* and *semantic context* associations over two phases: in the *generate* phase, they retrieved associations for a novel picture; in the *switch* phase they were shown the same picture and asked to retrieve a different association.

### 2.3. Materials

Stimuli were taken from the International Affective Picture System (IAPS; Lang et al., 2008), a database of pictures normed for valence and arousal. Thirty-six pictures were selected for *emotion* associations and were classified as either positive (valence mean > 6) or negative (valence mean < 4), with an equal number of positive and negative images in each run. Thirty-six pictures were selected for *semantic context* associations, all neutral valence (valence mean between 4 and 6). Content was broadly matched, such that *semantic context*, positive *emotion*, and negative *emotion* sets each contained equivalent proportions of people (61.1%) and animals (5.6%). Pictures for *semantic context* associations had significantly lower mean valence than positive *emotion* pictures (U < 1, *p* < .001), and higher mean valence than negative *emotion* pictures (U < 1, *p* < .001). Ratings of mean arousal were significantly higher for *emotion* than *semantic context* pictures (U = 221.0, *p* < .001), but were matched between positive and negative *emotion* images (U = 153.5, *p* = .791). Identifiers for each image and normed ratings of valence and arousal can be seen in Supplementary Table 1. Python scripts used to present them are available on the Open Science Framework (OSF; https://osf.io/498ur/).

### 2.4. Procedure

A summary of the procedure can be seen in Figure 1. Trials were presented across six functional runs lasting 4.5 minutes, each containing one *emotion* mini-block (six trials) and one *semantic context* mini-block (six trials). The order of trials within each run was consistent across participants. Run order was counterbalanced across participants such that a given block could appear in any position (first to sixth).

**Figure 1.**
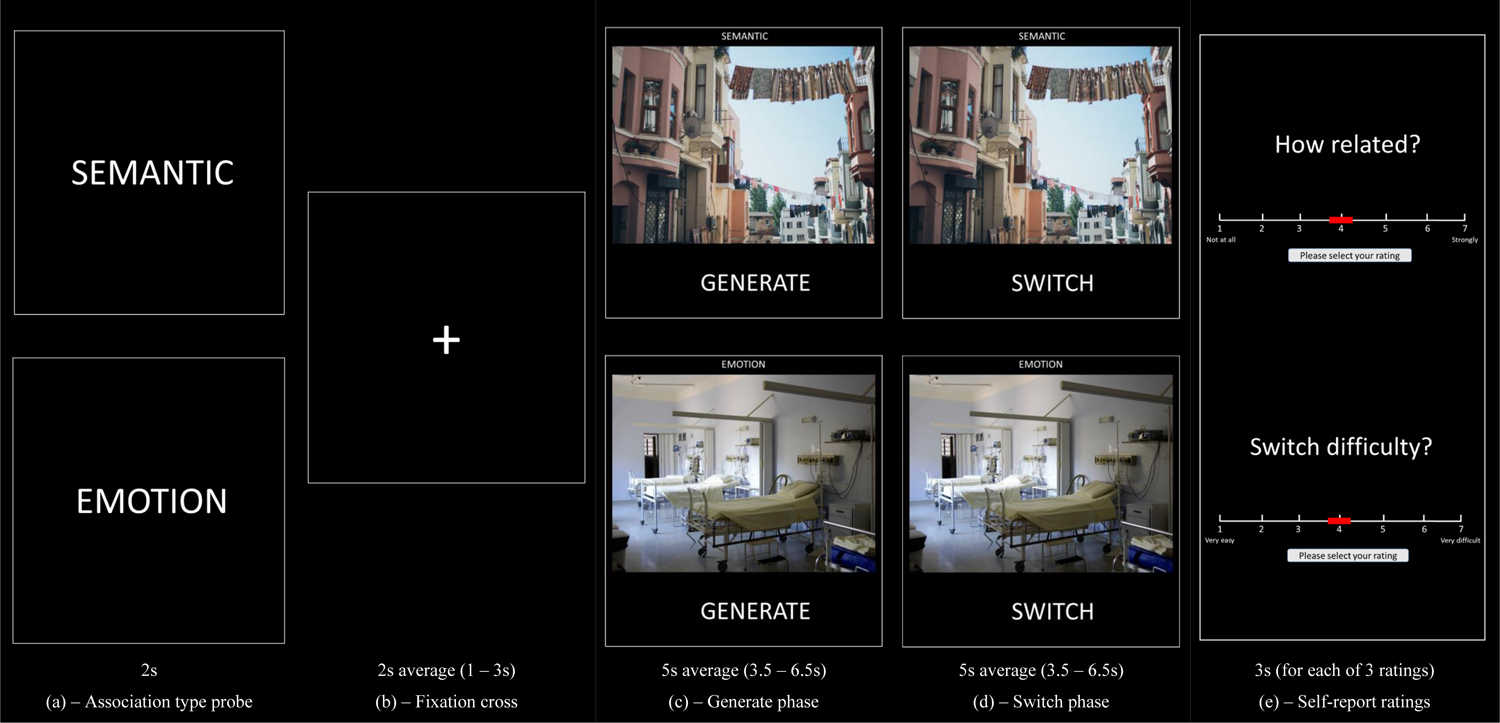
Examples of trials for both semantic context (top) and emotion (bottom) associations. Each miniblock was preceded by (a) a probe reflecting the current association type. Each trial was preceded by (b) a fixation cross. This was followed by (c) the ‘generate’ phase in which dominant response is generated and (d) the ‘switch’ phase in which a subordinate response is generated. Each trial contained (e) self-report ratings of strength of association between the picture and generated association after each phase, and overall switch difficulty at the end of the trial. The duration of each phase (in seconds) is indicated. Photos used for this figure are taken from stock photo website Pexels, from users Pixabay (emotion) and Elina Sazonova (semantic context)

The day before the scan, participants completed two practice runs (using stimuli not presented in the main experiment) remotely via Zoom (Zoom Video Communications Inc., 2016). During the first run participants reported their associations verbally, confirming they had correctly comprehended the study instructions. During the second run participants completed the task without verbal responses, to simulate the in-scanner procedure.

At the start of each mini-block, participants were presented with a 2s prompt informing them of the current association type (‘SEMANTIC’ or ‘EMOTION’). Each trial was preceded by a fixation cross, for an average of 2s, jittered between 1 and 3s. During the ‘GENERATE’ phase of *semantic context* associations, participants were shown a picture and asked to reflect on the first event-based association this picture brought to mind. Participants were told not to simply reflect on the content of the picture, and not to recall an episodic memory, but to identify a meaningful context from their general knowledge (e.g., for a picture of a street with washing lines, “visiting a city on holiday”). During the ‘SWITCH’ phase, participants were asked to stop reflecting on their initial association, and to reflect on another context associated with this picture (e.g., “using the sun to dry washing”). During the ‘GENERATE’ phase of *emotion* associations, participants were shown a picture and asked to reflect on how it made them feel. Participants were asked to avoid simple descriptive labels, but instead to embody emotions caused by the picture (e.g., for a picture of an empty hospital room, thinking about feeling sad). During the ‘SWITCH’ phase, participants were asked to stop thinking about this emotion, and to switch to another emotional response (e.g., fear).

Each *generate* and *switch* phase involved seeing a given picture for the first and second time, respectively. *Generate* and *switch* phases were presented for an average of 5s, jittered between 3.5 and 6.5s. After each *generate* and *switch* phase, participants indicated how strongly related their self-generated association was to the picture on a scale from 1 (no real relationship) to 7 (very strong relationship). At the end of each trial, participants indicated how difficult they found it to switch from the first to the second association on a scale from 1 (very easy) to 7 (very difficult). Each strength of association and difficulty rating was presented for a period of 3s.

Immediately following the scan, participants completed a recall assessment. Pictures were presented in the same order as in the scanner, and participants were asked to type the contexts or emotions they had generated, as well as their confidence in their recall accuracy (from 1-7), for both the *generate* and *switch* phases. This data was used to qualitatively validate that each participant had completed the task as intended. Another recent study focussed on the functional organisation of the anterior temporal cortex also includes an ROI-based analysis of this paradigm (Krieger-Redwood et al., 2023).

### 2.5. fMRI acquisition

Participants were scanned at the York Neuroimaging Centre, University of York, using a 3T Siemens MRI scanner with a 64-channel head coil, tuned to 123MHz. A localiser scan and six whole brain functional runs were acquired using a multi-band multi-echo (MBME) EPI sequence (TR = 1.5 s; TEs = 12, 24.83, 37.66 ms; 48 interleaved slices per volume with slice thickness of 3 mm (no slice gap); FoV = 24 cm (resolution matrix = 3×3×3; 80×80); 75° flip angle; 180 volumes per run; 7/8 partial Fourier encoding and GRAPPA (acceleration factor = 3, 36 ref. lines; multi-band acceleration factor = 2). Structural T1-weighted images were acquired using an MPRAGE sequence (TR = 2.3 s, TE = 2.26 s; voxel size = 1×1×1 isotropic; 176 slices; flip angle = 8°; FoV= 256 mm; interleaved slice ordering).

### 2.6. MRI data pre-processing

An MBME sequence was used to optimise signal from the medial temporal lobes while maintaining signal across the whole brain (Halai et al., 2014). Echoes within each functional run were combined using the TE Dependent ANAlysis (tedana; version 0.0.12; Kundu et al., 2011; 2013; the tedana community et al., 2021) library in Python. Pre-processing was performed before echoes were combined, using the Anatomical Processing Script pipeline in FSL (fsl_anat; https://fsl.fmrib.ox.ac.uk/fsl/fslwiki/fsl_anat). This included re-orientation to standard MNI space (fslreorient2std), automatic cropping (robustfov), bias-field correction (RF/B1 – inhomogeneity-correction, using FAST), linear and non-linear registration to standard-space (using FLIRT and FNIRT), brain extraction (using FNIRT, BET), tissue-type segmentation (using FAST) and subcortical structure segmentation (FAST). The multi-echo data were pre-processed using AFNI (https://afni.nimh.nih.gov/), including de-spiking (3dDespike), slice timing correction (3dTshift; heptic interpolation), and motion correction (with a cubic interpolation) of all echoes aligned to the first echo (3dvolreg applied to echo 1 to realign all images to the first volume; these transformation parameters were then applied to echoes 2 and 3).

### 2.7. Movement

To quantify movement during scanning, individual-level analyses were run on data corresponding to the second echo only, without motion correction and combination in tedana. Across the six runs, no participant presented with absolute mean displacement greater than .76mm (sample mean = .18mm), and no relative mean displacement greater than .17mm (sample mean = 0.06mm). No runs were excluded on the basis of movement.

### 2.8. fMRI data analysis

First-, individual-, and group-level analyses were conducted using FSL-FEAT version 6 (FMRIB’s Software Library, www.fmrib.ox.ac.uk/fsl; Jenkinson et al., 2012; Smith et al., 2004; Woolrich et al., 2009). Denoised optimally-combined time series output from tedana were submitted as input. Pre-processing included high-pass temporal filtering (Gaussian-weighted least-squares straight line fitting, with sigma = 50s), linear co-registration to native space using the respective participant’s structural T1-weighted image, and to MNI152 standard space (Jenkinson & Smith, 2001), spatial smoothing using a Gaussian kernel with full-width-half-maximum of 6 mm, and grand-mean intensity normalisation of the entire 4D dataset by a single multiplicative factor.

EVs in the model included time periods covering (1) *generate* and (2) *switch* phases of *emotion* associations, (3) *generate* and (4) *switch* phases of *semantic context* associations, (5) all self-report rating periods and (6) the association type prompt at the start of each miniblock. Fixation periods between trials were taken as the implicit baseline. Two parametric EVs reflected self-reported switch difficulty for (7) *semantic context* and (8) *emotion* associations, modelled for the *switch* phase of the respective trial. These parametric EVs included demeaned switch difficulty scores for each trial within a run, calculated separately for semantic and emotion associations. One run for one participant was excluded from the model as all *emotion* trials were rated the same for switch difficulty – resulting in each trial weighted as 0. Self-reported ratings of association strength for the *generate* and *switch* phase were not modelled as they both correlated with ratings of switch difficulty across the sample [*generate*: r_s_(2230) = −.14, *p* < .001, *switch*: r_s_(2234) = −.53, *p* < .001].

For whole-brain analysis at the group-level, we looked for activation associated with (1) either *semantic context* or *emotion* associations (across generate/switch phases) over baseline, as well as their conjunction (using FSL’s ‘eaythresh_conj’ tool), (2) contrasts of *generate* versus *switch* phases, (3) contrasts of *semantic context* versus *emotion* associations, (4) the interaction of phase and association type, and (5) parametrically higher or lower self-reported switch difficulty^3^. A threshold of Z > 3.1 was used for all group-level contrasts.

We characterised the placement of clusters associated with task activation on the principal gradient, reflecting the separation between unimodal and heteromodal regions. We performed Spearman spatial correlations between a map of the principal gradient (from Margulies et al., 2016) and unthresholded contrasts of each combination of association type and phase over baseline. This was performed at the individual-level, such that a coefficient was obtained for each contrast for each participant. We then ran a repeated-measures ANOVA examining effects and interactions of association type and phase. Positive mean coefficients reflect that a given condition falls towards the heteromodal end of the gradient, while negative coefficients show that conditions are towards the unimodal end. We focus here only on the principal gradient; analysis of gradients that explain the second and third most variance in connectivity can be seen in the Supplementary section ‘*Gradient 2 & 3 Analysis*’ (Supplementary Figure 3 and Supplementary Figure 4).

To search for differences between association types and to parse the function of DMN, we examined five resting-state networks taken from the 17-network parcellation from Yeo et al. (2011) that constitute 96% of voxels of the DMN resulting from the 7-network parcellation. These networks were (with % of the Yeo7-DMN in parentheses) ‘FT DMN’ (40.1%), ‘core DMN’ (37.8%), ‘control B’ (7.7%), ‘auditory’ (6.6%), and ‘MT DMN’ (3.8%). These network maps were mutually exclusive. The auditory and control B networks are often not considered to be DMN subnetworks but were included here as they constituted more of the Yeo7-DMN than the MT subnetwork, which is relatively small. We also assessed how much of each of these networks fell within the Yeo7-DMN: FT DMN = 95.3%, core DMN = 99.7%, auditory = 46.5%, MT DMN = 42.7%, control B = 23.4%. There is considerable variability in the size (number of voxels) of these networks: FT DMN = 12,672, core DMN = 11,435, auditory = 4,268, MT DMN = 2,680, control B = 9,951.

We ran region of interest (ROI) analysis using the Featquery function of FSL with binarised versions of the resting-state networks overlapping with the DMN. Full versions of DMN subnetworks were used as masks in ROI analysis (not only elements of these networks falling within the 7-network solution DMN). Mean percent signal change was calculated for each ROI in each combination of association type and phase over baseline. A repeated-measures ANOVA was run, examining effects and interactions of association type, phase, and network. There were two levels for association type (*semantic context, emotion*), two levels for phase (*generate, switch*), and five levels for network (FT DMN, core DMN, control B, auditory, FT DMN).

We also provide supplementary analyses (in section ‘*Functional control network analysis*’; Supplementary Tables 2-6 and Supplementary Figures 5-6) of functionally defined control networks, including SCN and MDN, as well as parts of these networks that overlap with each other and with DMN.

## 3. Results

### 3.1. Behavioural results

Descriptive statistics for self-reported association strength, switch difficulty, and recall confidence can be seen in Table 1. A repeated-measures ANOVA for self-reported association strength revealed significant main effects of association type (*emotion*/*semantic context*) [F(1, 31) = 17.1, *p* < .001, η_p_^2^ = .36] and phase (*generate*/*switch*) [F(1, 31) = 149.0, *p* < .001, η_p_^2^ = .83], but no association type by phase interaction [F(1, 31) = 1.6, *p* = .209, η_p_^2^ = .05]. These main effects reflect greater association strength in the *generate* than the *switch* phase, and for *emotion* than *semantic context* associations. No difference was found between association types (*emotion*/*semantic context*) for switch difficulty [t(31) = −1.8, *p* = .089]. For recall confidence, we observed a main effect of phase (*generate*/*switch*) [F(1, 31) = 80.5, *p* < .001, η_p_^2^ = .72], but no effect of association type (*emotion*/*semantic context*) [F(1, 31) = 0.8, *p*= .392, η_p_^2^ = .02] or association type by phase interaction [F(1, 31) < 0.1, *p* = .827, η_p_^2^ < .01]. Recall confidence was greater for the *generate* than the *switch* phase. These analyses confirm that any DMN activation for the *switch* versus *generate* phase is unlikely to reflect stronger associations being retrieved in this phase, or greater accessibility of memory when pictures are re-presented.

**Table 1.** Descriptive statistics for self-reported ratings of strength of association, switch difficulty, and recall confidence, split by association type.

|  | Mean (SD) |  |
| --- | --- | --- |
|  | Semantic context | Emotion |
| Generate association strength | 5.22 (0.69) | 5.60 (0.59) |
| Switch association strength | 4.00 (0.67) | 4.20 (0.62) |
| Switch difficulty | 4.05 (0.63) | 3.86 (0.64) |
| Generate recall confidence | 6.02 (0.58) | 5.94 (0.54) |
| Switch recall confidence | 5.26 (0.82) | 5.21 (0.78) |

Note: All ratings taken on a Likert scale from 1-7. Higher numbers reflect higher association strength, greater difficulty, and higher recall confidence.

To validate that participants were using an appropriate strategy for *semantic context* associations, we coded the content of all recalled responses across phases (see Supplementary Table 7). Most associations were ‘general semantic’ as instructed (70%). Some were classified as ‘personal semantic’ – associations reported in the first person and/or referring to a specific person or place in the participant’s life (7%). A small number were classified as ‘episodic’, as they alluded to discrete events in the participant’s life (3%). Given that these two latter categories still broadly conformed with the description of the *semantic context* condition (i.e., associating a context with the stimulus), these trials were not excluded from analysis. Although there was considerable variation across participants, *emotion* associations were largely recalled as single words [mean (SD) = 67.3% (27.4)].

### 3.2. Whole-brain analysis

Results of group-level whole-brain analysis are presented in Figure 2. Supplementary Figure 7 provides the percentage of voxels for each contrast that fall in each of the Yeo et al. (2011) 17 networks. Brain maps throughout this paper were visualised with the BrainNet Viewer (Xia et al., 2013; https://www.nitrc.org/projects/bnv/). Unthresholded versions of group-level NIFTI files for this project are available on Neurovault (https://neurovault.org/collections/CFYXAGAU/).

**Figure 2.**
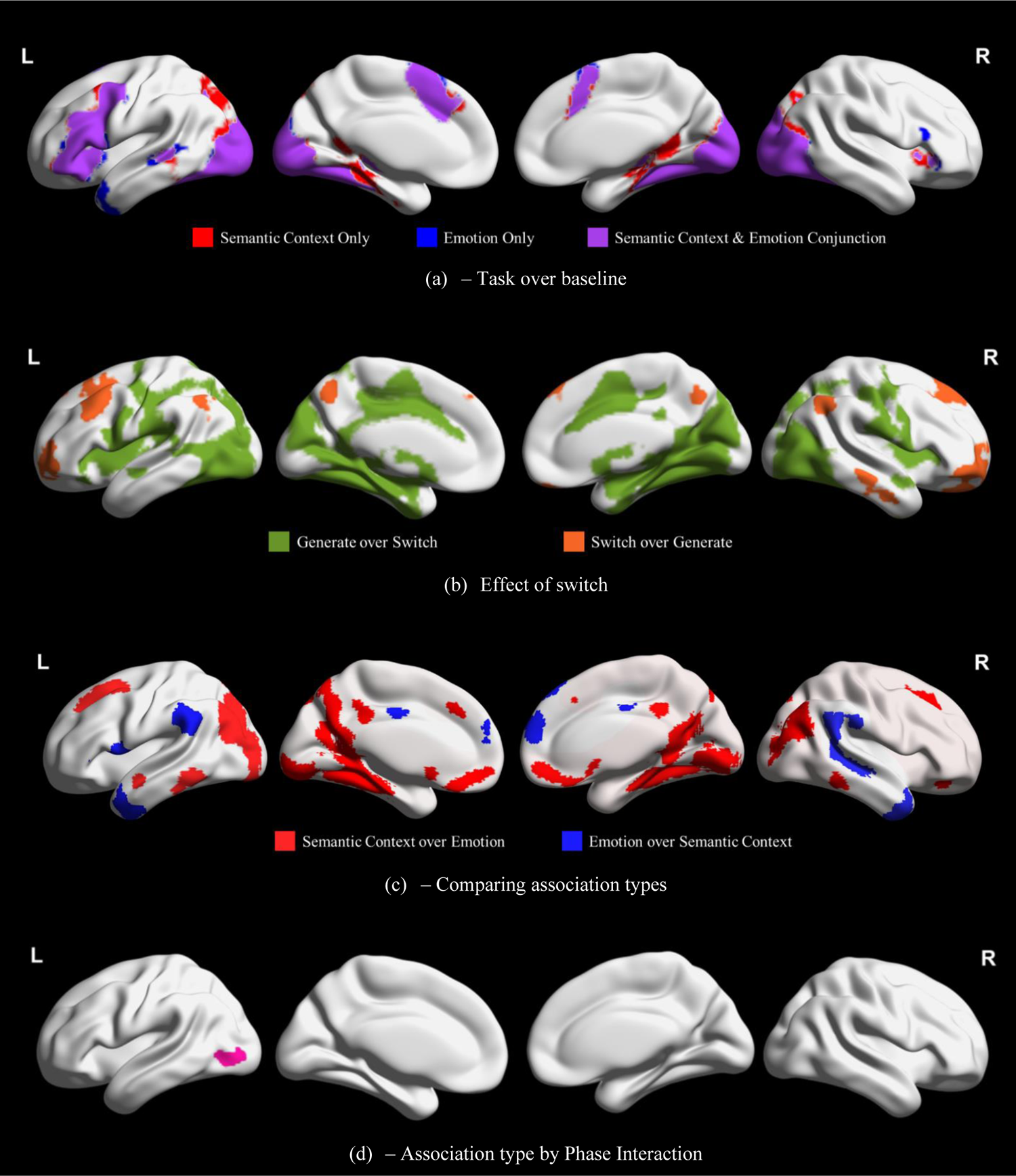
Clusters associated (a) with task activation relative to baseline, (b) more with one phase (generate or switch) than the other, (c) more with one association type (emotion or semantic context) than the other, and (d) in the interaction of association type and phase. Clusters taken from group-level analysis in FSL-FEAT with a threshold of Z > 3.1.

Figure 2a presents activation associated exclusively with either *semantic context* or *emotion* associations, as well as the conjunction of the two over baseline. Activation across association types is highly convergent. Key SCN clusters (Jackson, 2021) show activation across *emotion* and *semantic context* associations: (1) left IFG, middle frontal gyrus (MFG) and precentral gyrus, (2) left pMTG, (3) bilateral dorsomedial prefrontal cortex (dmPFC), (4) right IFG orbitalis; and for only emotion associations, (5) right IFG triangularis. Visual processing regions are also represented for both association types, including the occipital pole, lateral occipital cortex (LOC), and fusiform gyrus, as well as the bilateral thalamus and left caudate and pallidum.

Figure 2b presents clusters recruited more for the *switch* or *generate* phase, across association types. Clusters associated with the *generate* phase were extensive and highly overlapping with visual processing regions. The effect of *switch* fell within bilateral angular gyrus (AG), dmPFC, superior frontal gyrus (SFG), precuneus, and frontal pole, right pMTG and left MFG, with many of these clusters within DMN. As seen in Supplementary Figure 7e, this contrast shows the greatest overlap with the control B network (46.4%) followed by the core DMN (18.0%) and FT DMN (16.1%). We also found similarity in the effect of *switch* across association types, with overlapping effects across *emotion* and *semantic context* trials in bilateral frontal pole and dorsolateral PFC (see Supplementary Figure 8).

Figure 2c presents clusters recruited more for *semantic context* or *emotion* associations, across phases. Clusters associated with *semantic context* associations include bilateral LOC, occipital pole, pMTG, ventromedial PFC, posterior cingulate cortex (PCC), precuneus, MFG, SFG, and paracingulate gyrus, left anterior superior temporal gyrus, and right frontal pole. Much of the MT DMN (76%) is represented in these clusters, but this effect of association type extends to visual networks and DAN (given this broader recruitment, only 12.8% of the *semantic context* over *emotion* contrast falls within MT DMN; see Supplementary Figure 7b). In contrast, clusters showing greater activation for *emotion* associations include the bilateral temporal pole, supramarginal gyrus, dmPFC, and PCC, right AG and pMTG/posterior inferior temporal gyrus, and left IFG pars opercularis. This effect overlaps with the FT DMN; 11.5% of this DMN subnetwork falls within these clusters, with particular overlap in ATL, dmPFC, left IFG, and right pMTG. 34.4% of the *emotion* over *semantic context* contrast falls within FT DMN, with additional overlap in core DMN and language networks, as well as in ventral attention and limbic regions (see Supplementary Figure 7c).

Figure 2d presents the interaction of association type (*emotion*/*semantic context*) and phase (*generate*/*switch*), comprising one cluster in the left inferior LOC. We extracted mean percent signal change from the peak of this cluster in each phase and association type over baseline. There was a larger effect of phase for *emotion* associations (*generate* [.34; SD=.03] > *switch* [.18; SD=.03]), relative to *semantic context (generate* [.32; SD=.04] > *switch* [.21; SD=.03]) associations, in this cluster, primarily because the *switch* phase of *emotion* associations engaged this site less.

### 3.3. Gradient analysis

We next considered the distribution of task activation on the principal gradient of cortical organisation, which captures the distinction between heteromodal and unimodal cortex. We asked whether *emotion* and *semantic context* associations are located at different points in gradient space, and how the contrast of *generate* and *switch* phases for each of these association types changes this topographical pattern. Figure 3 provides visualisations of task contrasts from the whole-brain analysis above (‘*semantic context* over *emotion*’, ‘*emotion* over *semantic context*’, and ‘*switch* over *generate*’; Figure 3a), next to the principal gradient of intrinsic connectivity (Figure 3b), showing that all three of these effects overlap with the heteromodal end of the principal gradient (shown in red in Figure 3b), although the semantic context effect also extends into less heteromodal regions (shown in blue in Figure 3b).

**Figure 3.**
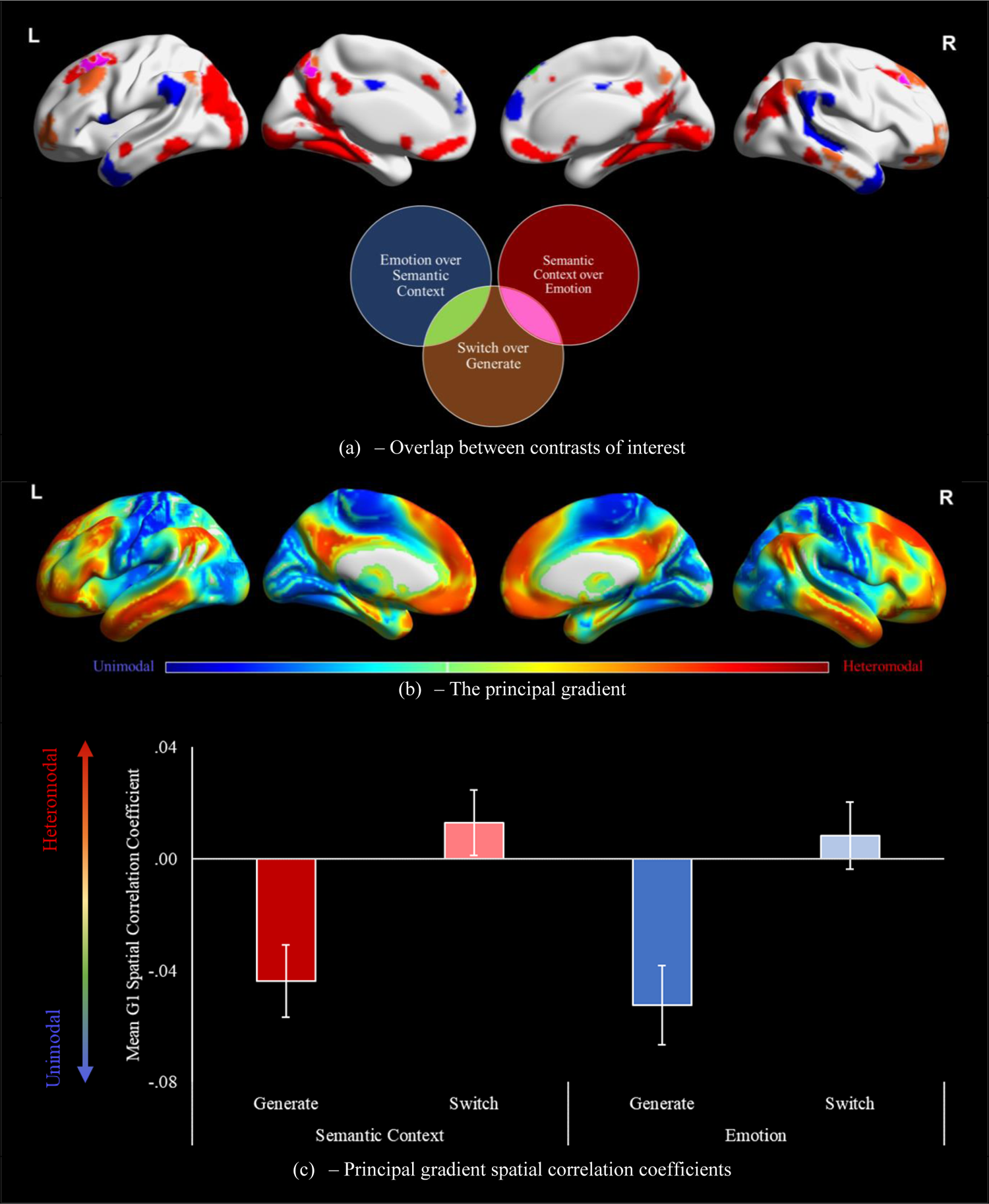
(a) Visualisations of overlap between the effect of switch and both contrasts of association type, and of (b) the principal gradient of cortical organisation. (c) Mean spatial correlation coefficients between the principal gradient and unthresholded contrasts of each association type and phase combination over implicit baseline. Error bars reflect one standard error. G1 = gradient 1.

Figure 3c presents the mean individual-level spatial correlation coefficients between the principal gradient and the unthresholded contrasts of each condition over baseline. A repeated-measures ANOVA of these spatial correlations revealed a significant main effect of phase [F(1, 31) = 58.9, *p* < .001, η_p_^2^= .66], but no effect of association type [F(1, 31) = 0.8, *p* = .36, η_p_^2^= .03] or interaction between these factors [F(1, 31) = 0.3, *p* = .599, η_p_^2^ = .01]. The main effect of phase reflected a stronger response towards the DMN-end of the gradient for the *switch* phase, and more sensory activation in the *generate* phase, consistent with our expectation that the *switch* phase would be less reliant on visual-to-semantic pathways and more reliant on heteromodal networks. Both association types showed this difference to the same degree: on the cortical surface this might correspond to a shift in the locus of activation away from DMN subnetworks linked to *semantic contexts* and *emotion* features, and towards common subnetworks across tasks. While the effects of association type were at equivalent positions on the gradient, clusters implicated specifically in the *switch* phase were more heteromodal. These findings demonstrate that heteromodal DMN regions, which lie towards the end of processing streams at a distance from unimodal cortex, show stronger engagement during memory retrieval when input cues are familiar. In contrast, regions supporting the retrieval of different aspects of knowledge (*semantic context* versus *emotion*), which are in partially distinct processing streams, fall at equivalent locations on these unimodal to heteromodal pathways. While we focus on the principal gradient of cortical organisation here, equivalent analysis of gradients that explain the second and third highest amount of variation can be seen in Supplementary Figure 3. The location of each DMN subnetwork on these three gradients can be seen in Supplementary Figure 4.

### 3.4. DMN overlap networks analysis

We next considered activation differences across phase and association type within DMN subnetworks, to establish the extent to which the effects of novelty of the retrieval cue and the type of knowledge being retrieved reflected previously-described functional dissociations within DMN. We used the Yeo et al. (2011) 7-network parcellation of intrinsic connectivity to identify voxels within DMN, broadly defined. We then used the Yeo 17-network parcellation to demarcate functional subdivisions within this DMN map, selecting five networks for analysis that showed maximum overlap with the Yeo-7 DMN. Figure 4a presents visualisations of the full networks used for analysis, that included voxels outside the Yeo-7 DMN. Figure 4b presents these same networks confined to the Yeo-7 DMN. We entered mean percent signal change for each network of interest into a repeated-measures ANOVA; data points above or below 3 standard deviations from the group mean were removed for each association type. All post-hoc tests were Bonferroni-corrected for five comparisons (reflecting five networks).

**Figure 4.**
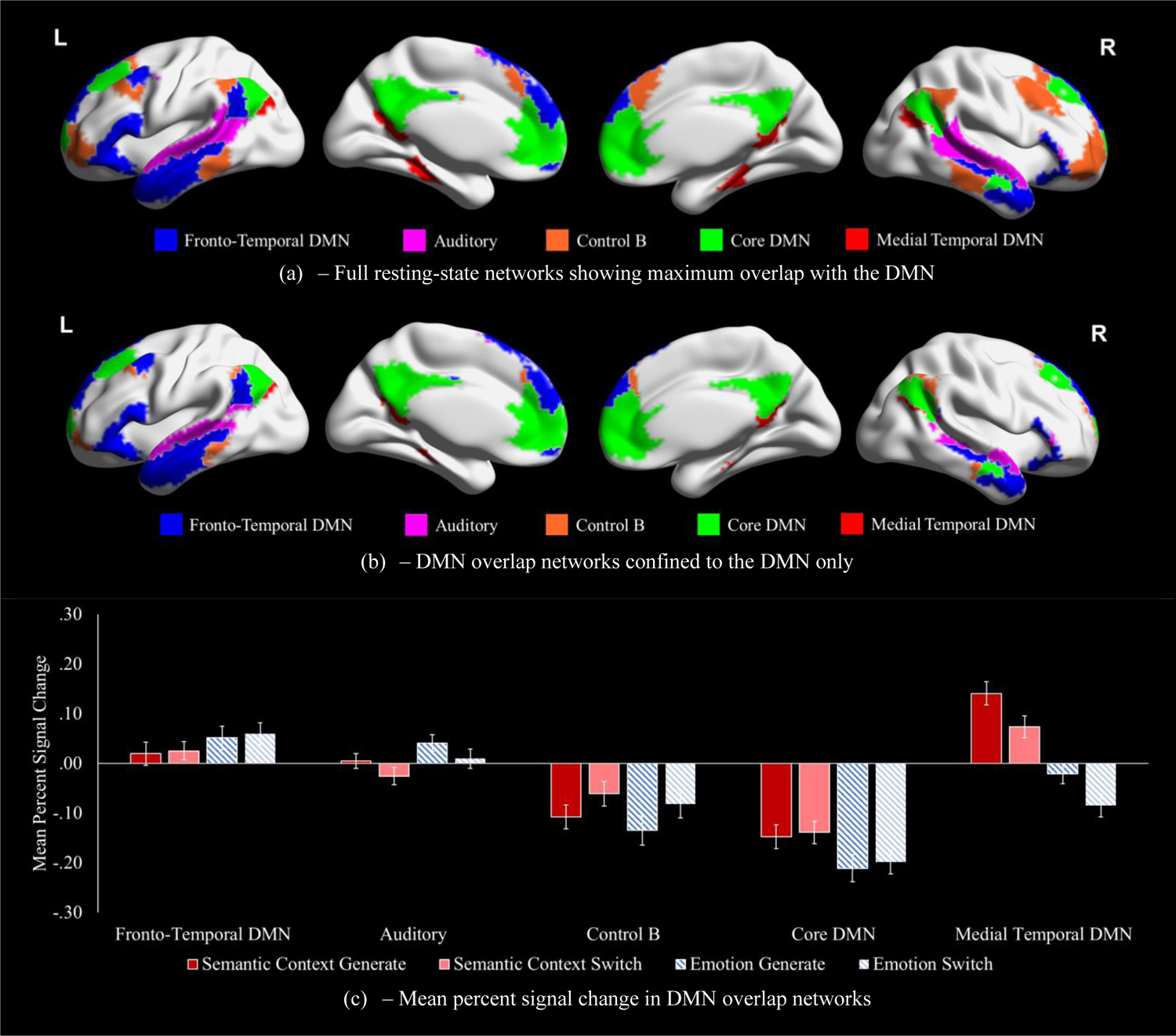
(a) Visualisation of complete resting-state networks showing maximum overlap with DMN, taken from the 17-network parcellation from Yeo et al., (2011) used to extract percentage signal change. (b) These networks restricted to voxels within the DMN from the 7-network parcellation. (c) Mean percent signal change in DMN resting-state networks in each combination of association type and phase, calculated using the Featquery function of FSL with binarised networks used as regions of interest. Error bars reflect one standard error. DMN = default mode network

Figure 4c presents mean percent signal change in each Yeo-17 network overlapping with DMN (shown in Figure 4a), for each condition over the implicit baseline. A repeated-measures ANOVA, reported in Table 2, revealed significant main effects of network, association type (*emotion*/*semantic context*), and phase (*generate*/*switch*), as well as an interaction of association type and network and of phase and network. The main effect of association type reflected, across all networks on average, higher mean percentage signal change for *semantic context* than *emotion,* while the phase main effect reflected higher mean percent signal change for *generate* than *switch*. Post-hoc comparisons examining the main effect of network are reported in Supplementary Table 8.

**Table 2.** Repeated measures ANOVA observing main effects and interactions of association type, phase, and network for mean percent signal change in resting-state networks overlapping with the default mode network.

| Analysis | Effect | Result |
| --- | --- | --- |
| Mean percent signal change | **Association type** | **$F(1, 28) = 52.1, p < .001, \eta_p^2 = .65^*$** |
|  | **Phase** | **$F(1, 28) = 6.3, p = .018, \eta_p^2 = .18^*$** |
|  | **Network** | **$F(1.4, 40.4) = 74.2, p < .001, \eta_p^2 = .73^*$** |
| | Association type x Phase | $F(1, 28) < 0.1, p = .778, \eta_p^2 < .01$ |
|  | **Association type x Network** | **$F(1.4, 40.0) = 69.4, p < .001, \eta_p^2 = .71^*$** |
|  | **Phase x Network** | **$F(1.6, 45.4) = 29.8, p < .001, \eta_p^2 = .52^*$** |
| | Association type x Phase x Network | $F(2, 56) = 0.5, p = .956, \eta_p^2 < .01$ |

Note: \* reflects a significant result. Assumption of sphericity violated for ‘network’. Greenhouse-Geisser adjustment applied accordingly.

For the network by association type interaction, MT DMN showed stronger activation for *semantic context* compared with *emotion* associations [t(30) = 11.4, *p* < .001]^4^, while core DMN showed less deactivation for the *semantic context* task [t(30) = 5.1, *p* < .001]. Two networks showed the reverse: more activation was seen for *emotion* than *semantic context* associations for FT DMN [t(30) = −2.9, *p* = .036] and the auditory network [t(30) = −4.3, *p =* .001]. These results show that DMN subsystems can capture the differences in activation relating to the retrieval of different aspects of knowledge (semantic context versus emotion), even though these responses were not in distinct positions on the principal gradient. The control B network did not differentiate between association types [t(30) = 1.6, *p* = .603].

For the network by phase interaction, more activation was seen in the *generate* than the *switch* phase for the MT DMN [t(30) = 5.8, p < .001]. The control B network showed less task-related deactivation for *switch* than *generate* [t(30) = −2.9, *p* = .035]. No difference was observed between phases for the core DMN [t(30) = −0.1, *p* > 1], FT DMN [t(30) = −1.5, *p* = .735], or auditory network [t(30) = 2.7, *p* = .060].

To summarise, MT and core DMN were linked to *semantic context* associations, while FT and auditory network showed stronger activation for *emotion* associations, consistent with recent accounts of functional specialisation in DMN (Andrews-Hanna and Grilli, 2021). The MT subnetwork also showed a preference for the *generate* phase, which likely required greater focus on visual features of the pictures, while control B (a control network allied to DMN) showed a preference for the *switch* phase, reflected by decreased deactivation. Control B responded equally to *semantic context* and *emotion* associations, suggesting that some DMN subsystems capture different processing streams from unimodal to heteromodal cortex, while others reflect location on these processing streams irrespective of whether contextual or evaluative information is retrieved.

This same analysis is provided for functional control networks, including SCN and MDN and their overlap with DMN in the supplementary materials (see Supplementary Tables 2-6 and Supplementary Figures 5-6). This analysis consistently implicated SCN, particularly within DMN, irrespective of association type or phase. MDN showed a preference for the generate phase and semantic context associations, potentially reflecting its greater proximity to sensorimotor cortex along the principal gradient (Wang et al., 2020).

### 3.5. Absence of task differences in the switch effect

A key research question is whether the contrasting retrieval demands of *generate* and *switch* phases differentially modulate the response of DMN subsystems linked to *semantic context* and *emotion* associations. In the analyses above, we observed no such differences in the recruitment of specific networks, subnetworks, or gradients. Nevertheless, in whole-brain analysis, we observed an interaction of association type and phase in left LOC. This implies that the mechanisms involved in generating associations from pictures as opposed to switching to new associations are largely orthogonal to the networks that process different association types, but with some overlap of these processes in higher-order visual regions. To stress-test this finding, we performed supplementary analysis. We split the activation highlighted in the thresholded *switch* over *generate* map into nine distinct, contiguous clusters (see Supplementary Table 9). We tested for differences within each cluster, using repeated measures ANOVA, reported in Supplementary Figure 9 and Supplementary Table 10. This analysis (see Supplementary Table 11) revealed that two clusters in this mask activated significantly more for *semantic context* than *emotion* associations. Despite this, no significant interaction was observed between association type and phase. We conclude that the effect of *switch* versus *generate* was largely not modulated by association type in heteromodal cortex. These effects were only recovered in LOC.

## 3. Discussion

This study advances our understanding of the functional organisation of cortical gradients, DMN, and control networks, by comparing activation during the retrieval of semantic context and emotional associations to pictures using fMRI. Each trial was split into a *generate* phase, thought to tap visual to DMN pathways, and a *switch* phase requiring a different association to be retrieved to the same picture, thereby increasing demands on internally-mediated retrieval processes. Accordingly, clusters identified in the *switch* phase were nearer the heteromodal end of the principal gradient than those in the *generate* phase, suggesting internally-oriented retrieval demands. A functional dissociation within DMN reflected the type of information needed in each task. *Semantic context* associations showed greater reliance on the MT subsystem, associated with scene construction. *Emotion* associations showed greater reliance on the FT subsystem, associated with abstract and evaluative processing. FT and MT subnetworks therefore showed a dissociation within semantic cognition when different meaning-based features were required. This was true regardless of retrieval demands, across the *generate* and *switch* phase. Across multiple analyses, we largely found similarities in this *switch* effect across association types, suggesting that this dimension of DMN organisation related to internal retrieval demands is largely orthogonal to the distinction between MT and FT subnetworks. Functional networks within DMN, including SCN, appear insensitive to the manipulation of association type, as demonstrated by supplementary analysis.

These findings contribute to our understanding of the functional specialisation of DMN subsystems. We found a dissociation between these networks when comparing tasks that tapped the retrieval of different meaning-based features (semantic contexts and emotions). This indicates that this subdivision relates to types of information processing; the MT subsystem supports the retrieval of general knowledge of meaningful contexts acquired over a lifetime. This is supported by studies implicating the MT subnetwork in contextually-specific and perceptually-guided scene construction and the FT subnetwork in abstract and evaluative processing (Andrews-Hanna et al., 2014; Andrews-Hanna and Grilli, 2021; Sheldon et al., 2019). Regions in FT DMN, including dmPFC, are involved in self-reflection about emotions and desires (Ochsner et al., 2004; van der Meer et al., 2010), suggesting the contribution of this network extends beyond simple semantic judgements. These results are consistent with the involvement of SCN, which shows considerable overlap with FT DMN in bilateral IFG, left pMTG, and dmPFC, in tasks requiring the regulation of emotional responses. Left IFG and pMTG have both been implicated in emotion reappraisal (Buhle et al., 2014; Kohn et al., 2014; Messina et al., 2015), a regulation strategy that relies on controlled processing (Braunstein et al., 2017). The IFG has also been associated with the suppression and substitution of emotional memories (Benoit & Anderson, 2012; Engen & Anderson, 2018; Guo et al., 2018). Overlapping elements of FT DMN and SCN may play a key role in the ability to interpret and reevaluate emotional categorisations.

The functional dissociation uncovered between semantic contexts and emotions in MT and FT also extended to additional DMN networks. The auditory network showed greater activation for *emotion* associations, while the core DMN showed less deactivation for *semantic context* associations. The Yeo et al. (2011) 17-network parcellation appears to be finer-grained than the functional dissociation recovered here, such that task differences extended over multiple linked networks. These network pairings may partly reflect spatial proximity – auditory and FT networks are adjacent in lateral temporal regions, while core and MT networks occupy adjacent positions in medial parietal cortex. In addition, language responses in the auditory network may be more relevant for emotion associations due to importance of language for abstract cognition. In any case, these observations support the claim that DMN is functionally organised according to the type of feature being processed.

We also observed a whole-brain interaction between association type and phase in left inferior LOC – implicated in visual processing of concepts (Coutanche & Thompson-Schill, 2015). The construction of rich scenes during *semantic context* associations may rely on this region across phases, while the *switch* phase of *emotion* associations may be abstract enough for decreased reliance. This mirrors the processing of abstract words, which show greater reliance on emotional content due to a relative lack of sensorimotor features (Kousta et al., 2011; Ponari et al., 2020; Rotaru & Vigliocco, 2019; Vigliocco et al., 2014). Gonzalez-Alam et al. (2018) found a similar region of inferior LOC to show an interaction between task demands and modality in the semantic domain, reflected by larger effects of inhibitory demands for picture-than word-based stimuli. The recruitment of visual object regions in controlled semantic retrieval appears to depend on both the nature of the input (modality) and the task demands (stimulus repetition).

In terms of effects of phase, resting-state network control B showed less task-related deactivation in the *switch* phase than the *generate* phase, suggesting it may play a role in the control of internally-oriented cognition. The *switch* effect was strongly overlapping with the control B network and located towards the DMN apex of the principal gradient, at a maximal distance from sensory-motor systems. The implication of transmodal regions in the switch phase is consistent with prior evidence. González-García et al. (2018) demonstrated that DMN regions towards the heteromodal end of the principal gradient support the representation of ambiguous Mooney images that have been disambiguated by participants upon second viewing. Our findings build on this by demonstrating the contribution of a specific resting-state network falling within DMN (control B), even when the stimuli themselves are not particularly ambiguous in nature (i.e., not actively degraded). Transmodal regions at the top of the principal gradient, and control B in particular, may play a role in reconfiguring representations of stimuli that have been previously encountered, shifting one’s reliance from perceptual properties to memory-guided processes.

Supplementary analysis suggested that SCN activation was not higher in the *switch* phase, despite evidence that both SCN and control B are allied to DMN (e.g., Davey et al., 2016; Dixon et al., 2018; Wang et al., 2020) and contribute to semantic cognition (e.g., Jefferies, 2013; Faber et al., 2019). This may be explained by the fact that SCN strongly overlaps with control A, with minimal overlap with control B (see Supplementary Figure 10). Control A is thought to support the control of externally-oriented cognition, as this network couples with dorsal attention regions to respond to external task demands (Yin et al., 2022). One reason for this distinction between the SCN and control B networks might be that SCN is defined according to activation in externally-presented tasks, while the *switch* phase of our experimental task did not involve the presentation of new stimuli. SCN might therefore support the controlled retrieval of semantic information from perceptual inputs as well as the internal generation of associations, while control B might be more critical for perceptually-decoupled semantic cognition.

Importantly, the observed functional dissociation in the FT and MT DMN subsystems in the processing of distinct semantic features was consistent across the *generate* and *switch* phases, with varying retrieval demands. Though the MT DMN did show a preference for the *generate* over the *switch* phase, this subnetwork still reliably activated for *semantic context* associations, while reliably deactivating for *emotion* associations. These subnetworks may therefore serve separate functions in a context-invariant manner, rather than themselves being sensitive to retrieval demands. Retrieval demands and task features may be largely orthogonal dimensions of DMN organisation. While showing a preference for the *switch* phase, control B showed no difference in activation between the *semantic context* and *emotion* tasks. Moreover, analysis positioning these task responses on whole-brain gradients capturing key dimensions of cortical organisation showed that clusters implicated in the *switch* phase tended towards the heteromodal apex of the principal gradient, while no effect of association type was observed. This suggests that *semantic context* associations, associated with the MT subnetwork, were not more perceptually-decoupled that *emotion* associations, associated with the FT subnetwork. This was true despite the MT subnetwork being associated with episodic memory (Andrews-Hanna & Grilli, 2021), even though many episodic memory tasks involve internally-oriented retrieval. Both MT and FT may be able to support access to heteromodal memory representations from visual inputs, as well as sustain more internal pathways to access spatial scenes and abstract, evaluative representations, thought to be supported by these subsystems, respectively.

There are limitations to the current study. First, the data do not indicate that the FT subsystem is not involved in the retrieval of semantic information about meaningful contexts; the analysis relies on task contrasts, so we can only conclude that the FT subsystem is less activated by *semantic context* than *emotion* associations. In this way, our data do not contradict the view that anterior and lateral temporal lobe regions act as a ‘semantic hub’, allowing us to integrate the full range of features that we learn about concepts (Lambon Ralph et al., 2017; Patterson et al., 2007). Second, the structure of our task does not disentangle the experience of seeing a picture for the first time from the generation of a dominant association. There are likely different levels of controlled processing required for the *generate* and *switch* phases, since participants indicated that their associations after the switch tended to be weaker. Future studies are needed to establish whether manipulations of the strength of the association being retrieved have comparable effects on MT and FT subnetworks, in the absence of any differences in the extent to which retrieval is externally- or internally-mediated. Third, while prior studies have implicated the MT DMN in episodic memory (Andrews-Hanna & Grilli, 2021), here we demonstrate its relevance in contextual semantic associations. We cannot say with certainty whether participants drew on episodic strategies in the retrieval of these associations, although participants were asked to avoid retrieving episodic memories. However, participants in prior work have reported using episodic strategies to generate strong semantic links between words (Krieger-Redwood et al., 2022). Future research on this topic may benefit from asking participants to provide detail on the strategy used to generate associations (e.g., Humphreys et al., 2022; Krieger-Redwood et al., 2022). Finally, it is unclear the extent to which participants were generating immersive visuospatial scenes and embodied emotional responses to stimuli. Findings from tasks requiring semantic judgements of valence, as conducted here, cannot be directly applied to experiential affect (Itkes & Kron, 2019). Future research may benefit from explicitly considering the role of semantic control in experiential aspects of semantic retrieval. For example, it is unclear if the same MT/FT dissociation would have occurred if the *emotion* associations had been more experiential in this study, given that the FT is associated with abstract aspects of cognition (Andrews-Hanna and Grilli, 2021).

## 4. Conclusion

We compared the neural mechanisms underlying the generation of both semantic contextual and emotional associations with pictures. Clusters implicated in the retrieval of subordinate-level associations were located towards the heteromodal end of the principal gradient, and showed reduced deactivation in a control network allied to DMN. The generation of contextual and emotional associations showed a dissociation across DMN subnetworks corresponding to scene construction and abstract processing, respectively. This dissociation was consistent across the *generate* and *switch* phases, suggesting that the functions of these networks are consistent despite varying retrieval demands.

## Acknowledgements

Thank you to Rebecca Lowndes, Richard Aveyard, Holly Brown, and Andre Gouws for their assistance in scanning participants and training investigators as MRI operators.

Thank you as well to Xiuyi Wang for her guidance with analysis.

## Footnotes

1 Sometimes referred to as the ‘dorsal medial subsystem’.

2 The ‘control A’ and ‘control B’ terminology is consistent with the labels frequently given to these respective sets of regions from the Yeo, et al. (2011) and Schaefer *et al*. (2018) 17-network parcellations. They have also often been referred to as ‘FPCNb’ and ‘FPCNa’, respectively.

3 Clusters corresponding to parametric analysis of difficulty can be seen in Supplementary Figure 1 and Supplementary Figure 2.

4 The assumption of normality was violated for tests of these interactions for both the control B and MT DMN networks. Non-parametric tests of these contrasts elicited the same outcomes. Association type comparison: control B [Z = −1.4, p = .763], MT DMN [Z = −4.9, p < .001]. Phase comparison: control B [Z = −2.7, p = .038], MT DMN [Z = −4.1, p < .001].

