## Supplementary Materials for "Default mode network shows distinct emotional and contextual responses yet common effects of retrieval demands across tasks"

Nicholas E. Souter [a, b], Antonia de Freitas [a], Meichao Zhang [a, c, d], Ximing Shao [a], Tirso Rene del Jesus Gonzalez Alam [a], Haakon Engen [e, f], Jonathan Smallwood [g], Katya Krieger-Redwood [a], Elizabeth Jefferies [a]

[a] Department of Psychology, University of York, York, UK

[b] School of Psychology, University of Sussex, Brighton, UK

[c] CAS Key Laboratory of Behavioral Science, Institute of Psychology, Beijing, China

[d] Department of Psychology, University of Chinese Academy of Sciences, Beijing, China

[e] Department of Psychology, University of Oslo, Oslo, Norway

[f] Institute for Military Psychiatry, Joint Medical Services, Norwegian Armed Forces, Norway

[g] Department of Psychology, Queen's University, Kingston, Ontario, Canada

**Corresponding Author**: Nicholas E. Souter, School of Psychology, University of Sussex, Falmer, BN1 9QH


Experimental Stimuli

Supplementary Table 1. Identifiers for stimuli taken from the International Affective Picture System with mean and SD of valence and arousal ratings, and allocation of association type and categorical valence in the current study.

| ID | Description | Valence Mean | Valence SD | Arousal Mean | Arousal SD | Association Type | Valence |
|---|---|---|---|---|---|---|---|
| 1120 | Snake | 3.79 | 1.93 | 6.93 | 1.68 | Emotion | Negative |
| 1350 | Pig | 5.25 | 1.96 | 4.37 | 1.76 | Semantic Context | Neutral |
| 1390 | Bees | 4.5 | 1.56 | 5.29 | 1.97 | Semantic Context | Neutral |
| 1659 | Gorilla | 6.57 | 1.98 | 4.89 | 1.97 | Emotion | Positive |
| 1675 | Buffalo | 5.24 | 1.48 | 4.37 | 2.15 | Semantic Context | Neutral |
| 1999 | Mickey | 7.43 | 1.47 | 4.77 | 2.4 | Emotion | Positive |
| 2002 | Man | 4.95 | 1.36 | 3.35 | 1.87 | Semantic Context | Neutral |
| 2036 | Woman | 5.8 | 1.28 | 3.24 | 1.88 | Semantic Context | Neutral |
| 2039 | Woman | 3.65 | 1.44 | 3.46 | 1.94 | Emotion | Negative |
| 2092 | Clowns | 6.28 | 1.9 | 4.32 | 2.29 | Emotion | Positive |
| 2156 | Family | 7.12 | 1.46 | 4.34 | 2.11 | Emotion | Positive |
| 2191 | Farmer | 5.3 | 1.62 | 3.61 | 2.14 | Semantic Context | Neutral |
| 2217 | Class | 6.24 | 1.52 | 4.08 | 1.85 | Emotion | Positive |
| 2360 | Family | 7.7 | 1.76 | 3.66 | 2.32 | Emotion | Positive |
| 2377 | Reading | 5.19 | 1.31 | 3.5 | 1.95 | Semantic Context | Neutral |
| 2382 | Artist | 5.67 | 1.19 | 3.75 | 1.97 | Semantic Context | Neutral |
| 2383 | Secretary | 4.72 | 1.36 | 3.41 | 1.83 | Semantic Context | Neutral |
| 2390 | Couple | 5.4 | 1.18 | 3.57 | 1.92 | Semantic Context | Neutral |
| 2397 | Men | 4.98 | 1.11 | 2.77 | 1.74 | Semantic Context | Neutral |
| 2455 | SadGirls | 2.96 | 1.79 | 4.46 | 2.12 | Emotion | Negative |
| 2456 | CryingFamily | 2.84 | 1.27 | 4.55 | 2.16 | Emotion | Negative |
| 2488 | Musician | 5.73 | 1.14 | 3.91 | 1.87 | Semantic Context | Neutral |
| 2489 | Musician | 5.66 | 1.44 | 3.8 | 1.93 | Semantic Context | Neutral |
| 2490 | Man | 3.32 | 1.82 | 3.95 | 2 | Emotion | Negative |
| 2595 | Women | 4.88 | 1.24 | 3.71 | 1.88 | Semantic Context | Neutral |
| 2635 | Cowboy | 5.22 | 1.65 | 4.42 | 1.98 | Semantic Context | Neutral |
| 2691 | Riot | 3.04 | 1.73 | 5.85 | 2.03 | Emotion | Negative |
| 2718 | DrugAddict | 3.65 | 1.58 | 4.46 | 2.03 | Emotion | Negative |
| 2745.1 | Shopping | 5.31 | 1.08 | 3.26 | 1.96 | Semantic Context | Neutral |
| 2751 | DrunkDriving | 2.67 | 1.87 | 5.18 | 2.39 | Emotion | Negative |
| 2870 | Teenager | 5.31 | 1.41 | 3.01 | 1.72 | Semantic Context | Neutral |
| 2980 | FoodBasket | 5.61 | 1.5 | 3.09 | 1.91 | Semantic Context | Neutral |
| 5300 | Galaxy | 6.91 | 1.8 | 4.36 | 2.62 | Emotion | Positive |
| 5455 | Cockpit | 5.79 | 1.37 | 4.56 | 2.17 | Semantic Context | Neutral |
| 5500 | Mushroom | 5.42 | 1.58 | 3 | 2.42 | Semantic Context | Neutral |
| 5621 | SkyDivers | 7.57 | 1.42 | 6.99 | 1.95 | Emotion | Positive |
| 5623 | Windsurfers | 7.19 | 1.44 | 5.67 | 2.32 | Emotion | Positive |
| 5814 | Mountain | 7.15 | 1.54 | 4.82 | 2.4 | Emotion | Positive |
| 5900 | Desert | 5.93 | 1.64 | 4.38 | 2.1 | Semantic Context | Neutral |

| | | | | | | | |
|---|---|---|---|---|---|---|---|
| 5910 | Fireworks | 7.8 | 1.23 | 5.59 | 2.55 | Emotion | Positive |
| 6240 | Gun | 3.79 | 1.8 | 5.27 | 2.2 | Emotion | Negative |
| 7001 | Buttons | 5.32 | 1.19 | 3.2 | 2.15 | Semantic Context | Neutral |
| 7033 | Train | 5.4 | 1.57 | 3.99 | 2.14 | Semantic Context | Neutral |
| 7036 | Shipyard | 4.88 | 1.08 | 3.32 | 2.04 | Semantic Context | Neutral |
| 7081 | Luggage | 5.36 | 1.3 | 3.96 | 2.24 | Semantic Context | Neutral |
| 7130 | Truck | 4.77 | 1.03 | 3.35 | 1.9 | Semantic Context | Neutral |
| 7234 | IroningBoard | 4.23 | 1.58 | 2.96 | 1.9 | Semantic Context | Neutral |
| 7325 | Watermelon | 7.06 | 1.65 | 3.55 | 2.07 | Emotion | Positive |
| 7492 | Ferry | 7.41 | 1.68 | 4.91 | 2.46 | Emotion | Positive |
| 7493 | Man | 5.35 | 1.34 | 3.39 | 2.08 | Semantic Context | Neutral |
| 7495 | Store | 5.9 | 1.6 | 3.82 | 2.33 | Semantic Context | Neutral |
| 7496 | Street | 5.92 | 1.66 | 4.84 | 1.99 | Semantic Context | Neutral |
| 7503 | CardDealer | 5.77 | 1.39 | 4.21 | 2.39 | Semantic Context | Neutral |
| 7506 | Casino | 5.34 | 1.46 | 4.25 | 1.95 | Semantic Context | Neutral |
| 7509 | Paintbrush | 6.03 | 1.35 | 3.43 | 2.02 | Emotion | Positive |
| 7520 | Hospital | 3.83 | 1.56 | 4.57 | 1.85 | Emotion | Negative |
| 7530 | House | 6.71 | 1.36 | 4 | 2.14 | Emotion | Positive |
| 7560 | Freeway | 4.47 | 1.65 | 5.24 | 2.03 | Semantic Context | Neutral |
| 7710 | Bed | 5.42 | 1.58 | 3.44 | 2.21 | Semantic Context | Neutral |
| 8158 | Hiker | 6.53 | 1.66 | 6.49 | 2.05 | Emotion | Positive |
| 8180 | CliffDivers | 7.12 | 1.88 | 6.59 | 2.12 | Emotion | Positive |
| 8312 | Golf | 5.37 | 1.41 | 3.32 | 2.06 | Semantic Context | Neutral |
| 8325 | RaceCars | 5.63 | 1.5 | 4.47 | 2.19 | Semantic Context | Neutral |
| 8499 | Rollercoaster | 7.63 | 1.41 | 6.07 | 2.31 | Emotion | Positive |
| 9090 | Exhaust | 3.56 | 1.5 | 3.97 | 2.12 | Emotion | Negative |
| 9110 | Puddle | 3.76 | 1.41 | 3.98 | 2.23 | Emotion | Negative |
| 9220 | Cemetery | 2.06 | 1.54 | 4 | 2.09 | Emotion | Negative |
| 9342 | Pollution | 2.85 | 1.41 | 4.49 | 1.88 | Emotion | Negative |
| 9445 | Skeleton | 3.87 | 1.57 | 4.49 | 2.01 | Emotion | Negative |
| 9622 | Jet | 3.1 | 1.9 | 6.26 | 1.98 | Emotion | Negative |
| 9630 | Bomb | 2.96 | 1.72 | 6.06 | 2.22 | Emotion | Negative |
| 9832 | Cigarettes | 2.94 | 1.58 | 4.46 | 2.06 | Emotion | Negative |

Parametric Difficulty Analysis

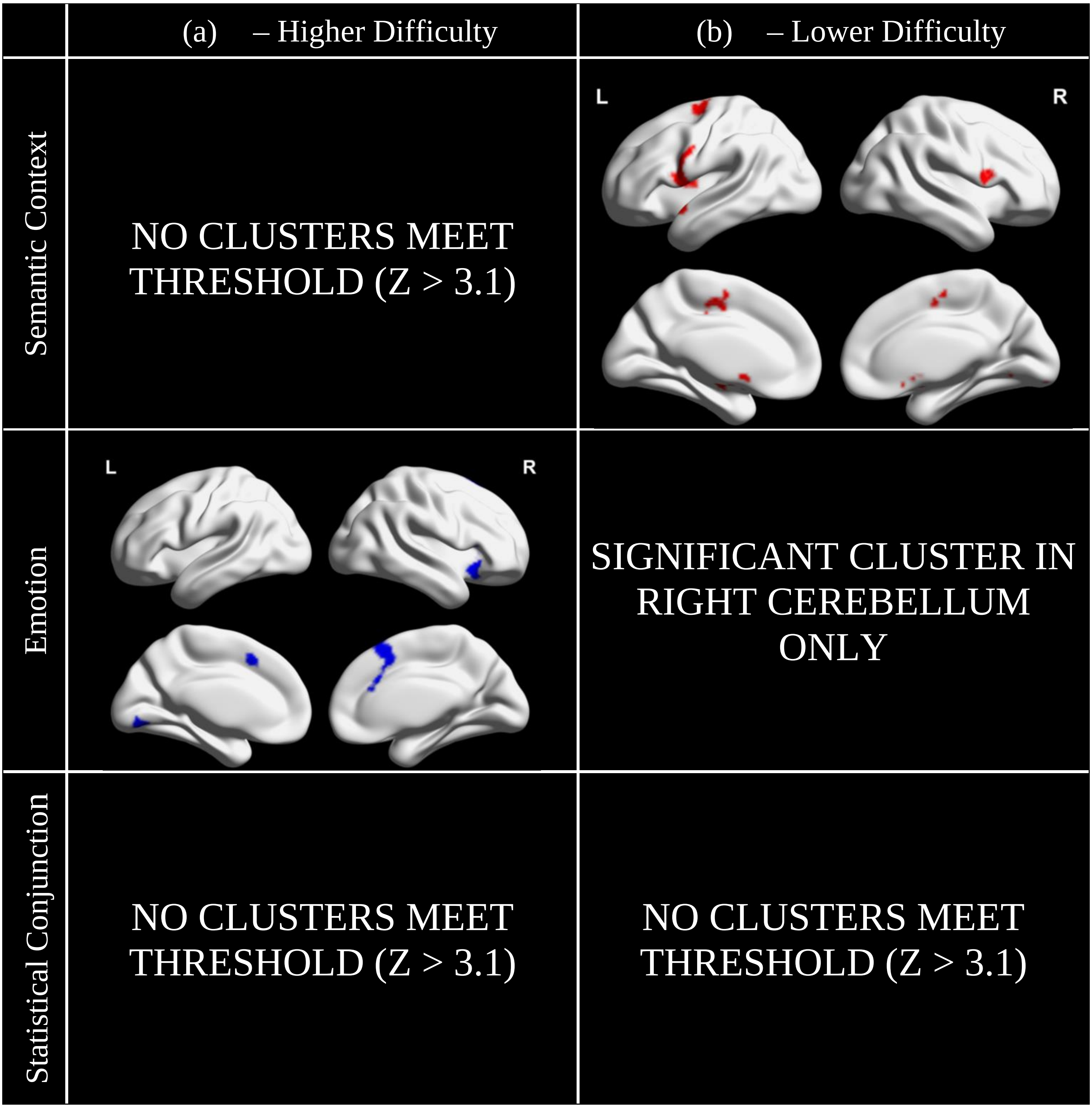


*Supplementary Figure 1. Clusters corresponding to (a) higher and (b) lower self-report switch difficulty during semantic context associations, emotion associations, and the statistical conjunction of the two. Clusters taken from group-level analysis in FSL-FEAT with a threshold of Z > 3.1. Maps visualised with the BrainNet Viewer (Xia et al., 2013; [https://www.nitrc.org/projects/bnv/](https://www.nitrc.org/projects/bnv/))*

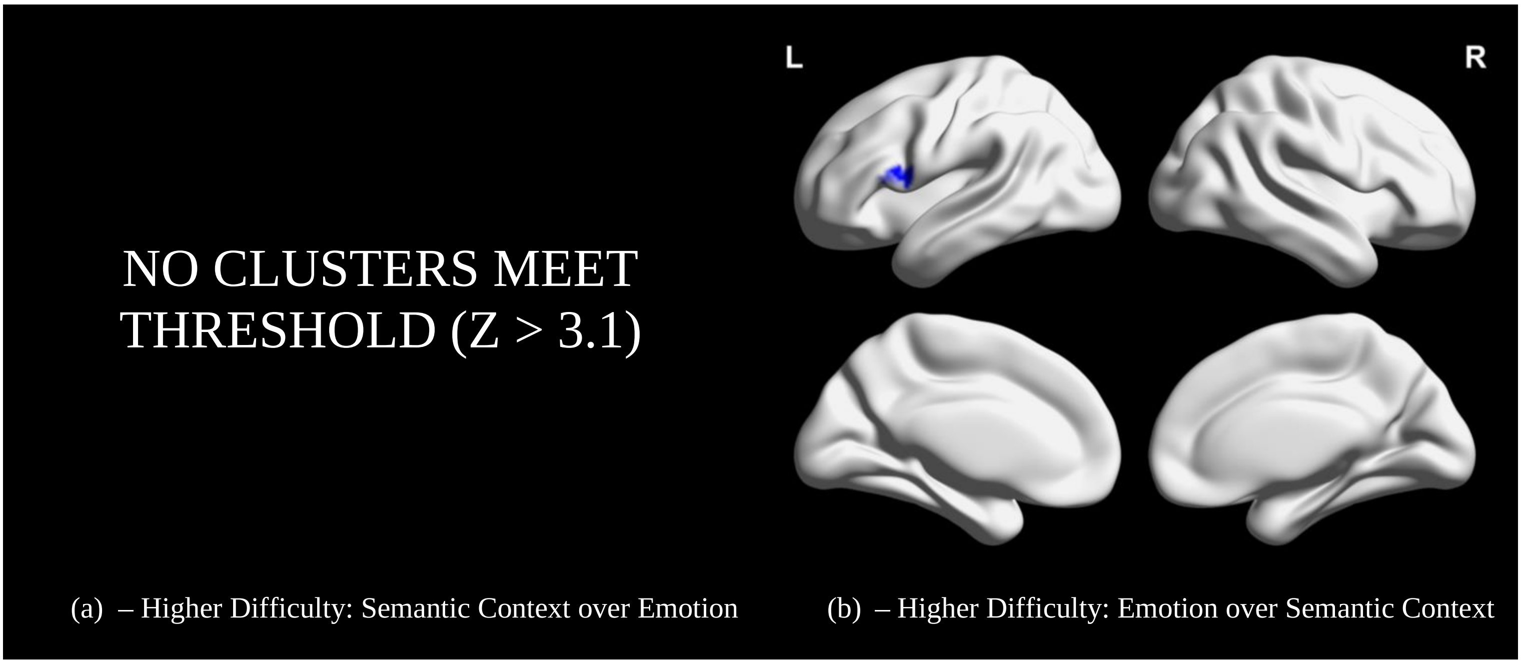


*Supplementary Figure 2. Clusters associated with (a) higher parametric self-reported difficulty for semantic context associations relative to emotion associations and (b) higher parametric self-reported difficulty for emotion associations relative to semantic context associations. Clusters taken from group-level analysis in FSL-FEAT with a threshold of Z > 3.1. Maps visualised with the BrainNet Viewer (Xia et al., 2013; [https://www.nitrc.org/projects/bnv/](https://www.nitrc.org/projects/bnv/))*

Interpretation

Supplementary Figure 1 displays clusters associated with both higher and lower self-reported switch difficulty, separately for both *semantic context* and *emotion* associations. The statistical conjunction across association types (generated using FSL's 'eaythresh_conj' tool) is also presented. For higher difficulty, clusters only survive thresholding for *emotion* associations. There are no significant clusters for *semantic context* associations or for the conjunction of association types. For lower difficulty, clusters for *semantic context* associations are observed in the cortex, with *emotion* associations only yielding one small cluster in the right cerebellum. Again, no clusters are implicated in the statistical conjunction. As seen in Supplementary Figure 2, there are no clusters associated more so with higher difficulty for *semantic context* associations compared to *emotion* associations. One cluster implicating aspects of the left inferior frontal gyrus and precentral gyrus is associated with greater difficulty for *emotion* associations, relative to *semantic context* associations.

### Gradients 2 & 3 Analysis

In the manuscript, we analysed individual level spatial correlations between the principal cortical gradient and unthresholded contrasts of each condition over baseline. Here, we present the same analysis for Gradient 2 (reflecting a separation between motor and visual regions) and Gradient 3 (reflecting a separation between DMN and frontoparietal regions; see Supplementary Figure 3). These gradients explain the second- and third-largest amounts of variance in the whole-brain decomposition of intrinsic connectivity by Margulies et al. (2016).

For gradient 2, we observed a main effect of association type [$F(1, 31) = 4.5$, $p = .041$, $\eta_p^2 = .13$] and of phase [$F(1, 31) = 18.4$, $p < .001$, $\eta_p^2 = .37$], but no association type by phase interaction [$F(1, 31) < 0.1$, $p = .981$, $\eta_p^2 < .01$]. The main effects reflect higher values (more visual responses) for *semantic context* associations than *emotion* associations, and for the *switch* phase compared with the *generate* phase. A stronger visual response for *semantic context* associations might reflect the relevance of visual-spatial scene construction processes in this condition. The main effect of phase is consistent with a greater relevance of motor information for the *generate* phase, compared to the *switch* phase.

For gradient 3, we observed a significant main effect of phase [$F(1, 31) = 7.2$, $p = .012$, $\eta_p^2 = .19$], but no main effect of association type [$F(1, 31) = 2.9$, $p = .097$, $\eta_p^2 = .09$] or association type by phase interaction [$F(1, 31) = 0.1$, $p = .764$, $\eta_p^2 < .01$]. The main effect of phase reflects greater reliance on frontoparietal control regions for the *switch* phase. Indeed, aspects of the control B network, implicated in clusters associated with the *switch* phase, sit further towards the frontoparietal control end of this gradient while other DMN-allied networks sit towards the lower end (see Supplementary Figure 4c).

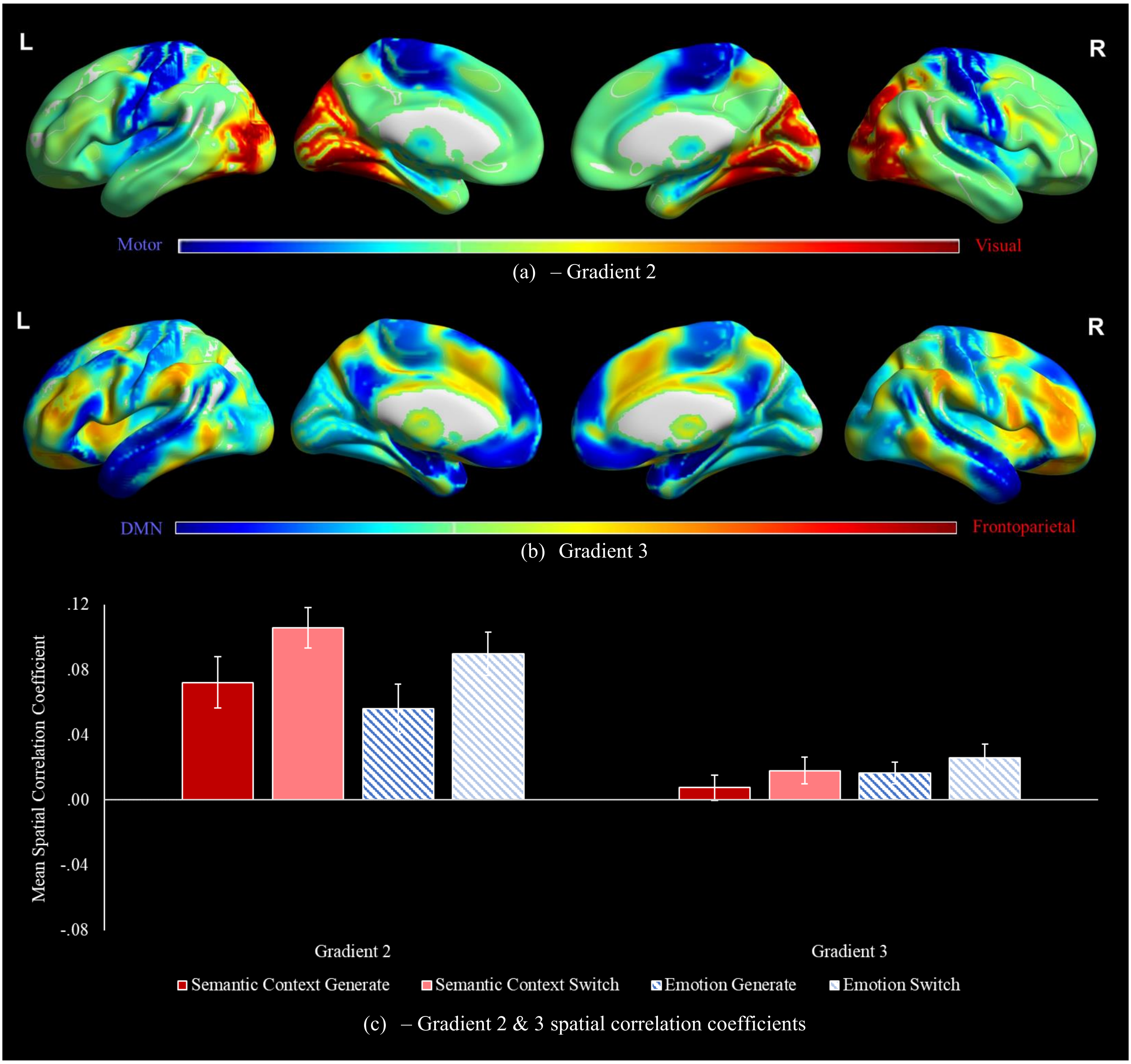


*Supplementary Figure 3. Visualisations of (a) gradient 2 and (b) gradient 3. (c) Mean spatial correlation coefficients between each gradient and unthresholded contrasts of each association type and phase combination over implicit baseline. Error bars reflect one standard error*

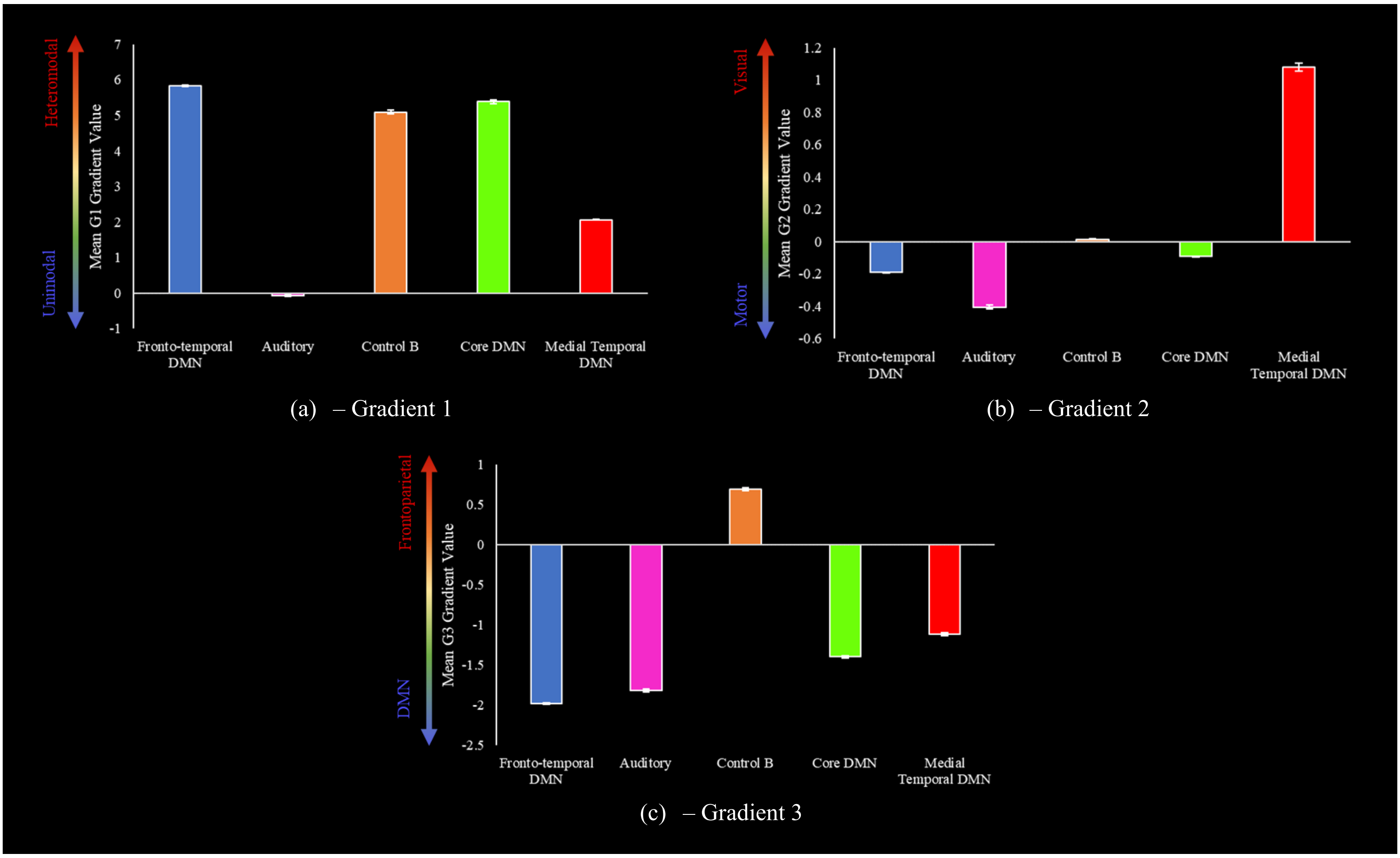


*Supplementary Figure 4. Mean gradient values for each resting-state DMN overlap network, for (a) gradient 1, (b) gradient 2, and (c) gradient 3. Error bars reflect one standard error. DMN = default mode network, G1 = gradient 1, G2 = gradient 2, G3 = gradient 3*

### Functional control network analysis

As for resting-state networks that overlap with DMN, we examined functional control networks. Control networks are organised systematically along the principal gradient, from DMN regions through semantic control regions to MDN regions implicated in domain-general control (Chiou et al., 2022; Wang et al., 2020). We therefore examined ROIs capturing this functional transition. SCN was defined using a meta-analysis of tasks with high > low semantic control demands (Jackson, 2021). A map of MDN was taken from Fedorenko et al. (2013), and reflects activation associated with more demanding versions of cognitive tasks, thresholded at t > 1.5. A map of DMN was taken from the 7-network parcellation from Yeo et al. (2011). All maps were mutually exclusive such that (1) voxels contained within both SCN and DMN were removed from each and placed in the 'DMN & SCN' map, and (2) any voxels contained within both SCN and MDN were removed from each and placed in the 'SCN & MDN'[1] map. This reduced the size of DMN by 6.4%, MDN by 8.0%, and SCN by 62.5%. These portions are captured in the joint overlap maps. The size (number of voxels) of the final network combinations varies: DMN = 28,191, DMN & SCN = 1,787, SCN = 2,389, SCN & MDN = 2,188, MDN = 26,457. Supplementary Table 2 provides the percentage of each network falling in each network from the Yeo et al. (2011) 17-network parcellation. Supplementary Figure 5a presents visualisations of these networks.

[1] 265 voxels overlapping in DMN and MDN were left in 'DMN'. 149 voxels overlapping in SCN, DMN, and MDN were left in 'SCN & MDN'. These areas of overlap account for small amounts of the overall networks (total voxel numbers; SCN = 6,364, MDN = 28,761, DMN = 30,127).

Supplementary Table 2. The percentage of each functional network (exclusive and conjunction) which falls within a given Yeo et al. (2011) 17-network parcellation.

| Network | Label | DMN | DMN & SCN | SCN | SCN & MDN | MDN |
|---|---|---|---|---|---|---|
| Yeo 1 | Visual peripheral | 0 | 0 | 0.29 | 0 | 8.77 |
| Yeo 2 | Visual central | 0 | 0 | 0 | 0 | 1.16 |
| Yeo 3 | Somatosensor A | 0 | 0 | 0 | 0 | 0.38 |
| Yeo 4 | Somatosensor B | 0 | 0 | 0 | 0 | 0.01 |
| Yeo 5 | DAN A | 0 | 0 | 4.94 | 9.28 | 13.19 |
| Yeo 6 | DAN B | 0 | 0 | 0.92 | 1.87 | 9.01 |
| Yeo 7 | VAN | 0 | 0 | 1.67 | 1.74 | 4.32 |
| Yeo 8 | Salience | 0.81 | 0.45 | 5.19 | 34.60 | 9.56 |
| Yeo 9 | Limbic A | 0 | 0 | 0 | 0 | 0 |
| Yeo 10 | Limbic B | 1.05 | 0.22 | 2.93 | 0 | 0 |
| Yeo 11 | Control C | 2.31 | 0 | 0 | 0 | 2.40 |
| Yeo 12 | Control A | 0 | 0 | 34.95 | 29.89 | 13.78 |
| Yeo 13 | Control B | 8.00 | 3.64 | 11.30 | 7.91 | 3.21 |
| Yeo 14 | Auditory | 6.62 | 6.66 | 2.22 | 0.50 | 0.07 |
| Yeo 15 | Medial temporal DMN | 4.06 | 0 | 0 | 0 | 0 |
| Yeo 16 | Core DMN | 40.39 | 0.45 | 0.08 | 0 | 0 |
| Yeo 17 | Fronto-temporal DMN | 36.77 | 88.58 | 5.90 | 7.82 | 0.04 |
| Total | | 100 | 100 | 70.41 | 93.60 | 65.89 |

Note: Totals for each network do not add to 100% in each case as aspects of some networks fall outside this 17-network parcellation. Colour grading used gives brighter colours to larger percentages. DAN = dorsal attention network, VAN = ventral attention network, DMN = default mode network, SCN = semantic control network, MDN = multiple demand network.

We ran region of interest (ROI) analysis using the Featquery function of FSL with binarised versions of these control networks. Mean percent signal change was calculated for each ROI in each combination of association type and phase over baseline. A repeated-measures ANOVA was run, examining effects and interactions of association type, phase, and network. There were two levels for association type (*semantic context, emotion*), two levels for phase (*generate, switch*), and five levels for network (DMN, DMN & SCN, SCN, SCN & MDN, MDN).

In our main analysis, a resting-state network associated with the control of memory – Control B – showed an effect of phase (a stronger response to familiar picture cues), yet no effect of type of information retrieved, this pattern may extend to functional control networks. Both the *generate* and *switch* phases are likely to draw on controlled retrieval but in different ways. The *generate* phase required participants to identify a specific association from an unfamiliar picture and selectively focus on this link (recruiting a pathway from visual cortex to DMN). In contrast, in the *switch* phase, the content of the picture has already activated memory systems that can drive a response, but it is necessary to inhibit the previous response and select a new one.

For all analyses below, for each network for each association type and phase, outliers (three standard deviations above or below the group mean) were excluded from analysis. All interaction post-hoc contrasts are Bonferroni-corrected for five comparisons (reflecting the five networks interrogated).

Supplementary Figure 5b presents mean percent signal change in each functional network, for each combination of association type and phase over implicit baseline. A repeated-measures ANOVA, shown in Supplementary Table 3, revealed a significant main effect of network, as well as interactions of association type (*emotion*/*semantic context*) and network, and of phase (*generate*/*switch*) and network. Post-hoc contrasts for the main effect of network are reported in Supplementary Table 4. Overall, the network capturing the overlapping regions of DMN & SCN showed more activation than all other networks (no difference was observed between the SCN and SCN & MDN, but both showed greater activation than both the DMN and MDN). Finally, MDN showed more activation than DMN. Post-hoc contrasts for the association type by network interaction revealed that MDN was recruited more for *semantic context* than *emotion* associations [t(31) = 2.9, $p$ = .038][2]. No difference between association types was observed for DMN [t(30) = 1.8, $p$ = .382], DMN & SCN [t(31) = -2.1, $p$ = .220], SCN [t(31) = 0.1, $p$ > 1], or SCN & MDN [t(31) = 2.0, $p$ = .265]. Post-hoc tests for the phase by network interaction revealed more activation in MDN during the *generate* than the *switch* phase [t(31) = 3.2, $p$ = .016]. No difference in phases was observed for DMN [t(30) = 0.2, $p$ > 1], DMN & SCN [t(31) = -0.5, $p$ > 1], SCN [t(31) = 0.5, $p$ > 1], or MDN & SCN [t(31) = 0.3, $p$ > 1][3].

[2] The assumption of normality was violated for the MDN in this post-hoc test – non-parametric testing of this effect did not survive correction [Z = -2.8, $p$ = .072].

[3] The assumption of normality was violated for the DMN, SCN, and SCN & MDN for this interaction. Non-parametric tests elicited the same outcomes: DMN [Z = -0.5, $p$ > 1], SCN [Z = -1.2, $p$ > 1], SCN & MDN [Z = -0.8, $p$ > 1].

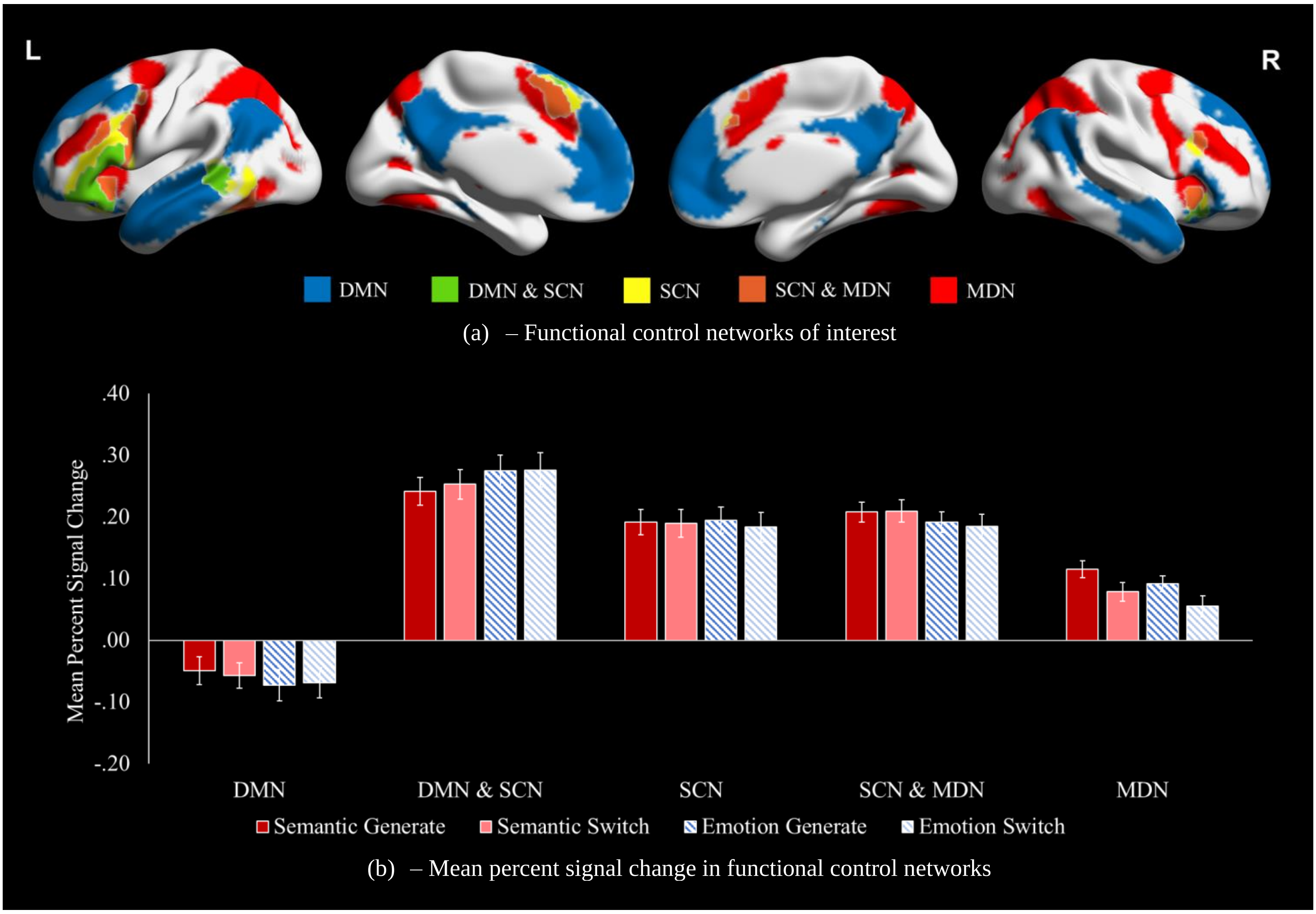


*Supplementary Figure 5. (a) Visualisation of functional control networks of interest. (b) Mean percent signal change in functional networks for each combination of association type and phase, calculated using Featquery in FSL, with binarised networks used as regions of interest. Error bars reflect one standard error. DMN = default mode network, SCN = semantic control network, MDN = multiple demand network*

Supplementary Table 3. Repeated measures ANOVA observing main effects and interactions of association type, phase, and network for mean percent signal change in functional control networks.

| Analysis | Effect | Result |
|---|---|---|
| Mean percent signal change | Association type | $F(1, 30) = 0.9$, $p = .338$, $\eta_p^2 = .03$ |
| | Phase | $F(1, 30) = 0.7$, $p = .398$, $\eta_p^2 = .02$ |
| | **Network** | **$F(2.9, 88.3) = 101.4$, $p < .001$, $\eta_p^2 = .77$*** |
| | Association type x Phase | $F(1, 30) < 0.1$, $p = .773$, $\eta_p^2 < .01$ |
| | **Association type x Network** | **$F(2.3, 68.0) = 6.5$, $p = .002$, $\eta_p^2 = .18$*** |
| | **Phase x Network** | **$F(2.0, 58.9) = 4.8$, $p = .012$, $\eta_p^2 = .14$*** |
| | Association type x Phase x Network | $F(2.4, 72.1) = 1.8$, $p = .160$, $\eta_p^2 = .06$ |

Note: * reflects a significant result. Assumption of sphericity violated for 'network'. Greenhouse-Geisser adjustment applied accordingly.

Supplementary Table 4. Post-hoc contrasts for the main effect of network on mean percent signal change in functional control networks.

| | DMN | DMN & SCN | SCN | SCN & MDN |
|---|---|---|---|---|
| DMN & SCN | **$t(30) = -18.9$, $p < .001$*** | - | - | - |
| SCN | **$t(30) = -15.6$, $p < .001$*** | **$t(31) = 5.8$, $p < .001$*** | - | - |
| SCN & MDN | **$t(30) = -12.5$, $p < .001$*** | **$t(31) = 3.7$, $p < .001$*** | $t(31) = -0.6$, $p > 1$ | - |
| MDN | **$t(30) = -7.8$, $p < .001$*** | **$t(31) = 8.1$, $p = .009$*** | **$t(31) = 5.6$, $p < .001$*** | **$t(31) = 7.7$, $p < .001$*** |

Note: post-hoc contrasts Bonferroni-corrected for ten comparisons. * reflects a significant result. DMN = default mode network, SCN = semantic control network, MDN = multiple demand network.

In summary, MDN showed a preference for the *generate* phase and *semantic context* associations, potentially reflecting its greater proximity to sensorimotor cortex along the principal gradient. Strong engagement of SCN was seen for all conditions, without differences between association types or phases; in this way, SCN did not resemble Control B.

To further parse the contribution of SCN, we conducted ROI analysis with the peaks of five distinct SCN clusters examining each condition over baseline (see Supplementary Figure 6 and Supplementary Table 5).

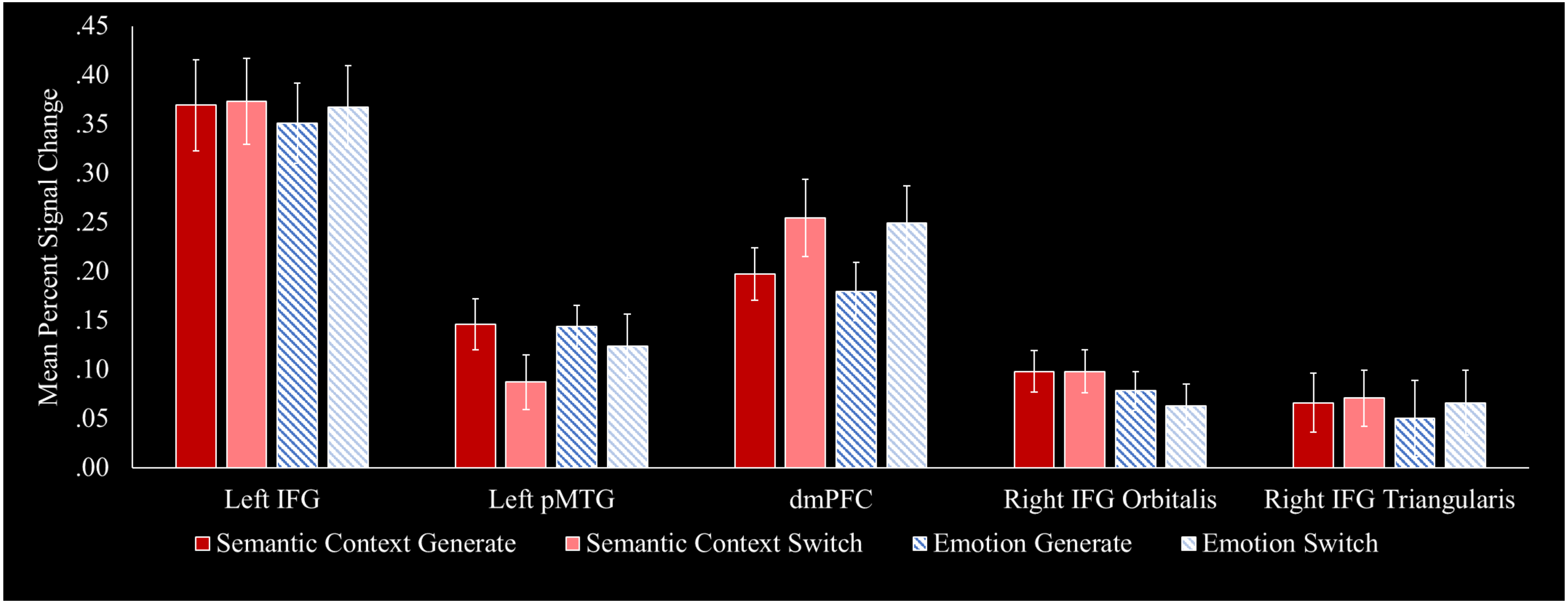


*Supplementary Figure 6. Mean percent signal change in the five peaks of the Jackson (2021) meta-analytics Semantic Control Network map, for each combination of association type (emotion or semantic context) and phase (generate or switch) over baseline. Error bars reflect one standard error. Analysis run using the Featquery function of FSL, with binarised 5mm spheres around each peak, based on voxel coordinates, used as regions of interest. IFG = inferior frontal gyrus, pMTG = posterior middle temporal gyrus, dmPFC = dorsomedial prefrontal cortex. N = 32*

Supplementary Table 5. A repeated measures ANOVA observing main effects and interactions of association type, phase, and network for the five peaks of the semantic control network.

| Effect | Result |
|---|---|
| Association type | F(1, 28) = 0.9, $p$ = .354, $\eta_p^2$ = .03 |
| Phase | F(1, 28) = 0.2, $p$ = .623, $\eta_p^2$ = .01 |
| **Peak** | **F(4, 112) = 19.9, $p$ < .001, $\eta_p^2$ = .42*** |
| Association type x Phase | F(1, 28) = 0.3, $p$ = .617, $\eta_p^2$ = .01 |
| Association type x Peak | F(4, 112) = 1.0, $p$ = .392, $\eta_p^2$ = .04 |
| **Phase x Peak** | **F(4, 112) = 5.3, $p$ < .001, $\eta_p^2$ = .16*** |
| Association type x Phase x Peak | F(2.8, 80.4) = 1.4, $p$ = .258, $\eta_p^2$ = .05 |

Note: * reflects a significant result.

The main effect of network is parsed below in Supplementary Table 6. These tests are Bonferroni-corrected for ten comparisons. This reveals significantly higher overall activation in the left IFG than in any other peak. In turn, the dmPFC shows higher overall activation than either right IFG cluster. No other differences between peaks were observed.

Supplementary Table 6. Post-hoc tests for the main effect of semantic control network peak on mean percent signal change.

| | Left IFG | Left pMTG | dmPFC | Right IFG orbitalis |
|---|---|---|---|---|
| Left IFG | - | - | - | - |
| Left pMTG | **t(29) = 6.5, $p$ < .001*** | - | - | - |
| dmPFC | **t(29) = 3.4, $p$ = .021*** | t(29) = -3.0, $p$ = .050 | - | - |
| Right IFG orbitalis | **t(30) = 6.5, $p$ < .001*** | t(30) = 0.9, $p$ > 1 | **t(30) = 3.6, $p$ = .010*** | - |
| Right IFG triangularis | **t(30) = 6.2, $p$ < .001*** | t(30) = 1.2, $p$ > 1 | **t(30) = 3.6, $p$ = .011*** | t(31) = 0.7, $p$ > 1 |

Note: * reflects a significant different. IFG = inferior frontal gyrus, pMTG = posterior middle temporal gyrus, dmPFC = dorsomedial prefrontal cortex.[4]

Post-hoc comparisons for the phase by peak interaction are Bonferroni-corrected for five comparisons. While no contrast survives correction, the left pMTG trends towards being more implicated in the *generate* phase than the *switch* phase [t(30) = 2.7, $p$ = .061]. Conversely, the dmPFC trends towards being more implicated in the *switch* phase than the *generate* phase [t(30) = -2.5, $p$ = .083]. Highly non-significant differences between phases was observed for the left IFG [t(30) = -0.7, $p$ > 1], right IFG orbitalis [t(31) = 1.0, $p$ > 1], and right IFG triangularis [t(31) = -0.5, $p$ > 1].

[4] Assumption of normality violated for the pMTG and dmPFC. However, non-parametric tests elicit the same results: pMTG – left IFG [Z = -4.5, $p$ < .001], pMTG – dmPFC [Z = -2.6, $p$ = .104], pMTG – right IFG orbitalis [Z = -1.1, $p$ > 1], pMTG – right IFG triangularis [Z = -1.1, $p$ > 1], dmPFC – left IFG [Z = -2.9, $p$ = .041], dmPFC – right IFG orbitalis [Z = -3.3, $p$ = .011], dmPFC – right IFG triangularis [Z = -3.3, $p$ = .009].

Interpretation of semantic control network results

Despite the observed functional dissociation in the engagement of DMN subsystems, the *emotion* and *semantic context* tasks showed overlapping activation in SCN nodes including left IFG, left pMTG, bilateral dmPFC, and right IFG. This suggests that the contribution of SCN cuts across distinct aspects of memory-based cognition that rely on the FT and MT subnetworks. Previous studies have reported an important role for SCN in contextual associations (Krieger-Redwood et al., 2022). The contribution of this network to emotion associations is predicted by the view that valence is a semantic feature (Martin, 2016; Lambon Ralph et al., 2017) and that the capacity to access emotion categories (e.g., fear) relies on effective semantic cognition (Lindquist et al., 2015). The representation of affective similarity correlates with the overall associative similarity of abstract words in left ATL (Meersmans et al., 2020), and valence congruency facilitates the retrieval of thematic associations between words (Marino Dávolos et al., 2020). Post-stroke impairments in semantic control are also associated with impaired categorisation of facial portrayals of emotion (Souter et al., 2021) and classification of words by valence (Souter et al., 2023). Key SCN sites, including left IFG and pMTG, have been implicated in emotion reappraisal (Buhle et al., 2014; Kohn et al., 2014; Messina et al., 2015), an emotion regulation strategy that relies on controlled processing (Braunstein et al., 2017). The IFG has also been associated with the suppression and substitution of emotional memories (Benoit & Anderson, 2012; Engen & Anderson, 2018; Guo et al., 2018). Together, our findings suggest this network plays a role in the control of tasks that tap internal representations, including the retrieval of emotion features and meaning-based scenes. Indeed, the contribution of SCN may extend beyond semantic processing given that this network has been implicated in other domains including episodic retrieval (Cogdell-Brooke et al., 2020; Stampacchia et al., 2018; Vatansever et al., 2021) and social cognition (Binney & Ramsey, 2020; Diveica et al., 2021).

Overall, SCN supported semantic retrieval processes in *generate* and *switch* phases equally; however, analysis of individual SCN peaks revealed greater importance of left pMTG in the *generate* phase, and of dmPFC in the *switch* phase. Left pMTG has been shown to support semantic control across verbal and picture modalities (Krieger-Redwood et al., 2015; Jackson, 2021). Given its proximity to visual and auditory input regions and the focus of the *generate* phase on the initial retrieval of semantic information from an external cue, this finding suggests a stronger role of pMTG in externally-mediated semantic cognition

(Davey et al., 2016). In contrast, dmPFC might make a greater contribution to controlling internally-mediated patterns of retrieval or in supporting the cognitive switching process. Activation in dmPFC has previously been associated with the generation of unusual links between word pairs (Krieger-Redwood et al., 2022), consistent with the demands of the *switch* phase in the current study. We also found that MDN showed a modest preference for the *generate* phase. This might reflect the fact that MDN encompasses frontoparietal and dorsal attention networks and is therefore well-suited to supporting external control.

Coding Semantic Context Recall Responses

Supplementary Table 7. The mean percentage (standard deviation) of semantic context associations that fall in each coding category, split by phase and overall.

| | General semantic | No association made | Personal semantic | Picture label | Episodic | Emotion association retrieved | Can't remember |
|---|---|---|---|---|---|---|---|
| Generate | 75.8 (20.3) | 3.8 (5.8) | 8.9 (10.7) | 4.6 (8.6) | 2.9 (5.4) | 0.3 (1.1) | 3.8 (7.1) |
| Switch | 64.7 (18.1) | 10.9 (10.8) | 5.6 (8.2) | 1.9 (4.2) | 3.3 (7.6) | 0.4 (1.0) | 13.2 (14.5) |
| Overall | 70.2 (17.7) | 7.3 (7.1) | 7.3 (8.6) | 3.3 (6.1) | 3.1 (6.3) | 0.4 (0.9) | 8.5 (10.1) |

All categories are mutually exclusive, such that percentages add to 100% in each phase. Scores are taken from coding of recall data. It is not possible to say how accurately these reflect associations made in the scanner, although participants were generally confident in their recall (see Table 1). Criteria used for each category are explained below:

- **General semantic** – Appropriate *semantic context* associations that reflect contextual associations free from personal semantics or specific episodes.
- **No association made** – Trials where participant report not having been able to generate an association for a given phase.
- **Personal semantic** – Associations that are semantic in nature but refer to specific people or places in the participant's life, and/or are reported in the first person.
- **Picture label** – Brief descriptions which do not provide anything beyond the content of the picture, e.g., "gambling" for a picture of a casino.
- **Episodic** – Associations that appear to reflect a specific event or memory in the participant's life, suggesting episodic recall rather than generation of a semantic contextual association.
- **Emotion association retrieved** – Associations for which participants appear to have mistakenly thought they were in an *emotion* block, and as such have reported an emotional response to a *semantic context* stimulus.
- **Can't remember** – Participants report not being able to remember the association they generated.

<u>Yeo Network Overlap</u>

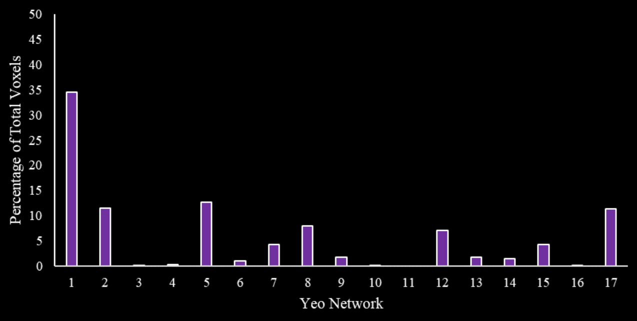


(a) – Task over Baseline

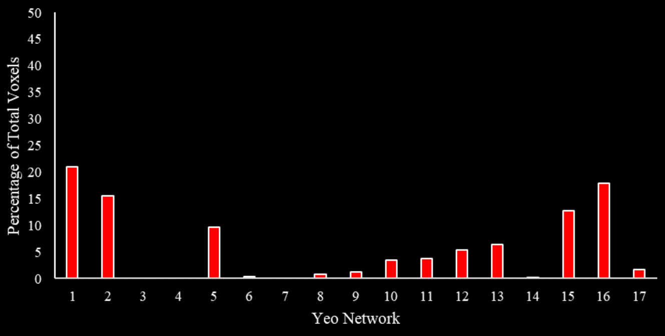


(b) – Semantic Context over Emotion

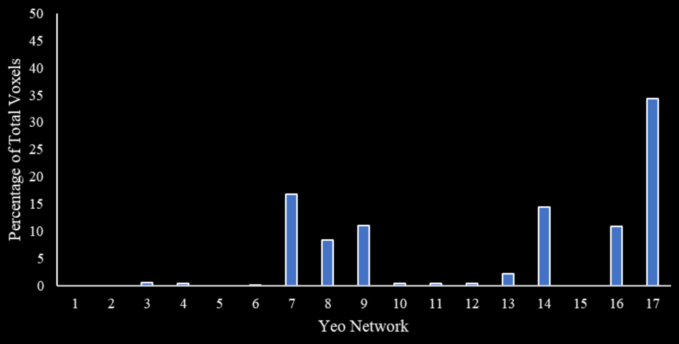


(c) – Emotion over Semantic Context

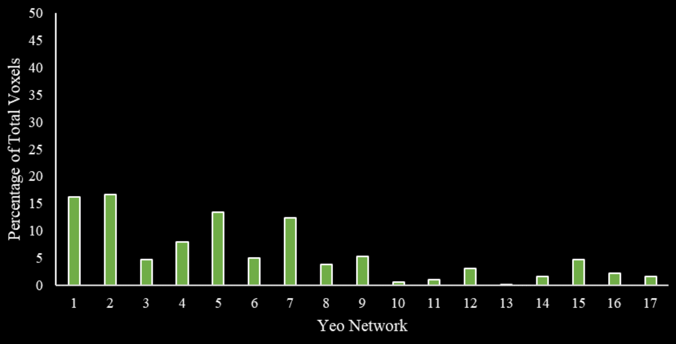


(d) – Generate over Switch

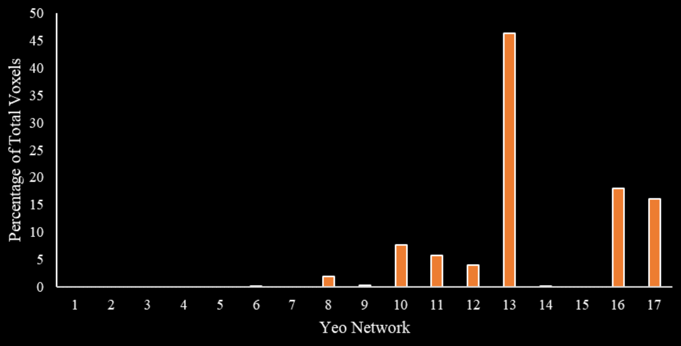


(e) – Switch over Generate

| | | | | | |
|---|---|---|---|---|---|
| Yeo 1 | Visual peripheral | Yeo 2 | Visual central | Yeo 3 | Somatosensor A |
| Yeo 4 | Somatosensor B | Yeo 5 | DAN A | Yeo 6 | DAN B |
| Yeo 7 | VAN | Yeo 8 | Salience | Yeo 9 | Limbic A |
| Yeo 10 | Limbic B | Yeo 11 | Control C | Yeo 12 | Control A |
| Yeo 13 | Control B | Yeo 14 | Auditory | Yeo 15 | Default C (Medial Temporal) |
| Yeo 16 | Default A (Core) | Yeo 17 | Default B (Fronto-Temporal) | | |

(f) – Yeo et al. (2011) 17-network parcellation labels

*Supplementary Figure 7. The percentage implicated in each of the Yeo et al. (2011) 17-network parcellation for clusters associated with (a) the conjunction of both semantic context and emotion associations over baseline, (b) greater activation for semantic context than emotion associations, (c) greater activation for emotion than semantic context associations, (d) greater activation in the generate than the switch phase, and (e) greater activation in the switch than the generate phase. Each contrast was masked by the addition of all 17 networks for this figure, such that bars for each contrast add up to 100% together*[5]*. (f) Labels are presented for each network.*

[5] Variable portions of each contrast fell within the addition of all 17 Yeo networks before masking: task over baseline [70.0%], *semantic context* over *emotion* [85.5%], *emotion* over *semantic context* [89.5%], *generate* over *switch* [62.3%], *switch* over *generate* [93.5%].

### Switch over Generate Split by Condition

Supplementary Figure 8 shows the effect of *switch* over *generate* separately for the *emotion* and *semantic context* conditions, and the overlap of these effects. The threshold of Z > 3.1 shows some clusters that only show an effect of *switch* in the emotion condition (Supplementary Figure 8a). The data have therefore also been presented at a lower threshold of Z > 2.6, demonstrating that clusters are similar across conditions, despite these effects being weaker for *semantic context* associations.

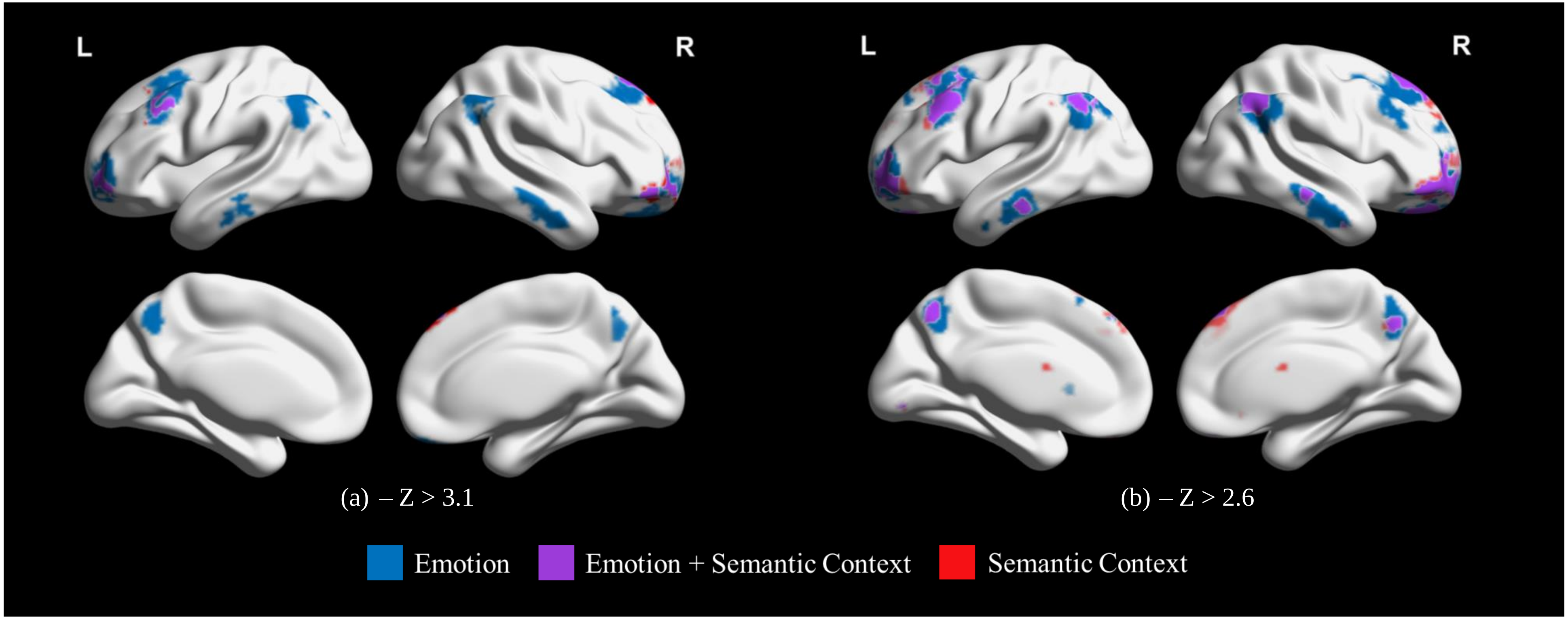


*Supplementary Figure 8. Clusters associated more with the switch than the generate phase split by association type (emotion or semantic context) at thresholds of (a) Z > 3.1 and (b) Z > 2.6. Maps visualised with the BrainNet Viewer (Xia et al., 2013;* [https://www.nitrc.org/projects/bnv/](https://www.nitrc.org/projects/bnv/)*)*

<u>DMN Subnetwork Mean Percent Signal Change Network Main Effect Contrasts</u>

Supplementary Table 8. Post-hoc contrasts for the main effect of network on mean percent signal change in resting-state networks overlapping with the DMN.

| | Control B | Auditory | Medial temporal DMN | Core DMN |
|---|---|---|---|---|
| Auditory | **t(29) = -5.3, *p* < .001*** | - | - | - |
| Medial temporal DMN | **t(29) = -4.3, *p* = .002*** | t(29) = -1.2, *p* > 1 | - | - |
| Core DMN | **t(29) = 4.2, *p* = .002*** | **t(29) = 10.5, *p* < .001*** | **t(29) = 8.8, *p* < .001*** | - |
| Fronto-temporal DMN | **t(29) = -6.9, *p* < .001*** | t(29) = -2.2, *p* = .327 | t(29) = -0.6, *p* > 1 | **t(29) = -18.5, *p* < .001*** |

Note: post-hoc contrasts Bonferroni-corrected for ten comparisons. * reflects a significant result. DMN = default mode network.[6]

[6] The assumption of normality was violated for both the control B network and medial temporal DMN. While parametric contrasts are used in this table, non-parametric equivalents elicited the same outcomes: control B – auditory [Z = -3.9, *p* = .001], control B – medial temporal DMN [Z = -3.5, *p* = .004], control B – core DMN [Z = -3.5, *p* = .004], control B – fronto-temporal DMN [Z = -4.6, *p* < .001], medial temporal DMN – auditory network [Z = -0.7, *p* > 1], medial temporal DMN – core DMN [Z = -4.8, *p* < .001*], medial temporal DMN – fronto-temporal DMN [Z = -0.5, *p* > 1].

Analysis of Contiguous Clusters within the Switch over Generate Contrast

To test whether effects of association type on the effect of *switch* were being masked, we separated the thresholded contrast of *switch* over *generate* into all its separate contiguous clusters. There were nine clusters in total. The location and size of these are characterised in Supplementary Table 9.

Supplementary Table 9. Contiguous clusters that comprise the thresholded (Z > 3.1) switch over generate contrast, including characterisation of regions implicated, size in voxels, and location in MNI coordinates.

| | Cluster Number | | | | | | | | |
|---|---|---|---|---|---|---|---|---|---|
| | 1 | 2 | 3 | 4 | 5 | 6 | 7 | 8 | 9 |
| Regions | R cerebellum | L AG<br>L pSMG<br>L superior LOC | R AG<br>R pSMG<br>R superior LOC | Precuneus | R pMTG<br>R aMTG<br>R pSTG<br>R pITG | L frontal pole | R SFG<br>R MFG<br>R frontal pole<br>L SFG<br>L frontal pole | L SFG<br>L MFG<br>L frontal pole | R frontal pole<br>R OFC |
| Size (voxels) | 222 | 237 | 345 | 360 | 480 | 626 | 772 | 1099 | 1104 |
| Peak (MNI) | (42, -68, -45) | (-40, -70, 44) | (48, -54, 48) | (-6, -62, 50) | (58, -12, -27) | (-30, 62, -2) | (12, 32, 54) | (-38, 20, 44) | (15, 41, -23) |

Note: R = right, L = left, AG = angular gyrus, pSMG = posterior supramarginal gyrus, LOC = lateral occipital cortex, pMTG = posterior middle temporal gyrus, aMTG = anterior middle temporal gyrus, pSTG = posterior superior temporal gyrus, pITG = posterior inferior temporal gyrus, SFG = superior frontal gyrus, MFG = middle frontal gyrus, OFC = orbitofrontal cortex.

We used binarised versions of each cluster as ROIs in Featquery, interrogating them relative to each combination of association type and phase over implicit baseline. Doing so may reveal whether any of these clusters were particularly important for either association type. Outliers were removed at each level by identifying data points three standard deviations above or below the group mean. The resulting data can be seen in Supplementary Figure 9. Results of a repeated measures ANOVA run on this data can be seen in Supplementary Table 10.

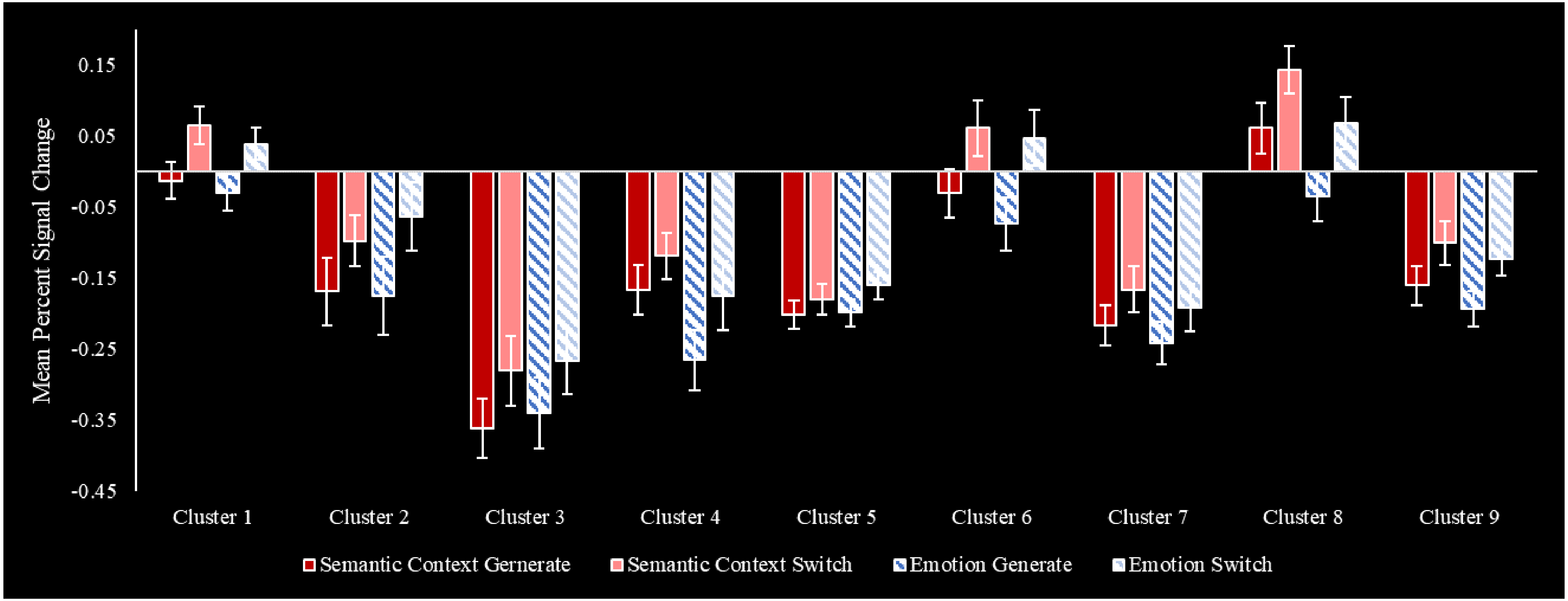


*Supplementary Figure 9. Mean percent signal change in contiguous clusters that comprise the overall thresholded (Z > 3.1) switch over generate contrast for each combination of association type and phase over implicit baseline. Error bars reflect one standard error. N = 32*

Supplementary Table 10. Repeated measures ANOVA for effects of association type and phase on contiguous switch over generate clusters.

| Effect | Result |
|---|---|
| Association type | **F(1, 29) = 4.6, *p* = .041, $\eta_p^2$ = .14*** |
| Phase | **F(1, 29) = 25.6, *p* < .001, $\eta_p^2$ = .47*** |
| Cluster | **F(5.3, 153.6) = 30.6, *p* < .001, $\eta_p^2$ = .51*** |
| Association type by Phase | F(1, 29) = 2.3, *p* = .137, $\eta_p^2$ = .08 |
| Association type by Cluster | **F(2.5, 73.0) = 4.2, *p* = .013, $\eta_p^2$ = .13*** |
| Phase by Cluster | **F(4.7, 135.3) = 4.3, *p* = .002, $\eta_p^2$ = .13*** |
| Association type by Phase by Cluster | F(4.9, 142.2) = 1.2, *p* = .296, $\eta_p^2$ = .04 |

Note: * reflects a significant effect. Assumption of sphericity violated for ‘cluster’. Greenhouse-Geisser adjustment applied accordingly.

The significant effect of association type reflects more overall activation for *semantic context* than *emotion* associations, while the main effect of phase reflects more overall activation for the *switch* phase than the *generate* phase. The main effect of cluster was not of interest here, and as such post-hoc tests of this effect are not reported. Post-hoc contrasts for the observed interactions can be seen in Supplementary Table 11. As seen here, two clusters activated significantly more for the *semantic context* than the *emotion* condition. Unsurprisingly, given that these clusters came from the *switch* over *generate* contrast, all clusters except two activate significantly more for the *switch* than *generate* phase (although the two which do not meet significance after correction are marginal). These results do imply a slight preference of *switch* clusters for *semantic context* associations. However, the absence of any interaction between association type and phase here is consistent with overall evidence of parity in *switch* effects across association types.

Supplementary Table 11. Post-hoc contrasts for significant switch over generate cluster interactions.

| Cluster | Association type by cluster | Phase by cluster |
|---|---|---|
| 1 | t(31) = 1.5, *p* > 1 | **t(31) = -4.9, *p* < .001*** |
| 2 | t(31) = -0.3, *p* > 1[7] | **t(31) = -4.2, *p* = .002*** |
| 3 | t(30) = -0.9, *p* > 1 | t(30) = -2.8, *p* = .086 |
| 4 | **t(30) = 4.8, *p* < .001*** | **t(30) = -3.8, *p* = .006*** |
| 5 | t(31) = -0.9, *p* > 1 | t(31) = -2.9, *p* = .055 |
| 6 | t(31) = 1.3, *p* > 1 | **t(31) = -5.2, *p* < .001***[8] |
| 7 | t(31) = 1.5, *p* > 1 | **t(31) = -3.2, *p* = .029*** |
| 8 | **t(31) = 5.8, *p* < .001*** | **t(31) = -5.5, *p* < .001*** |
| 9 | t(31) = 2.1, *p* = .435 | **t(31) = -5.4, *p* < .001*** |

Note: * reflects a significant difference.

[7] Assumption of normality violated, non-parametric test elicits the same result: Z = -3.7, *p* = .002*

[8] Assumption of normality violated, non-parametric test elicits the same result: Z = -4.2, *p* < .001*

### SCN and Control Network Overlap

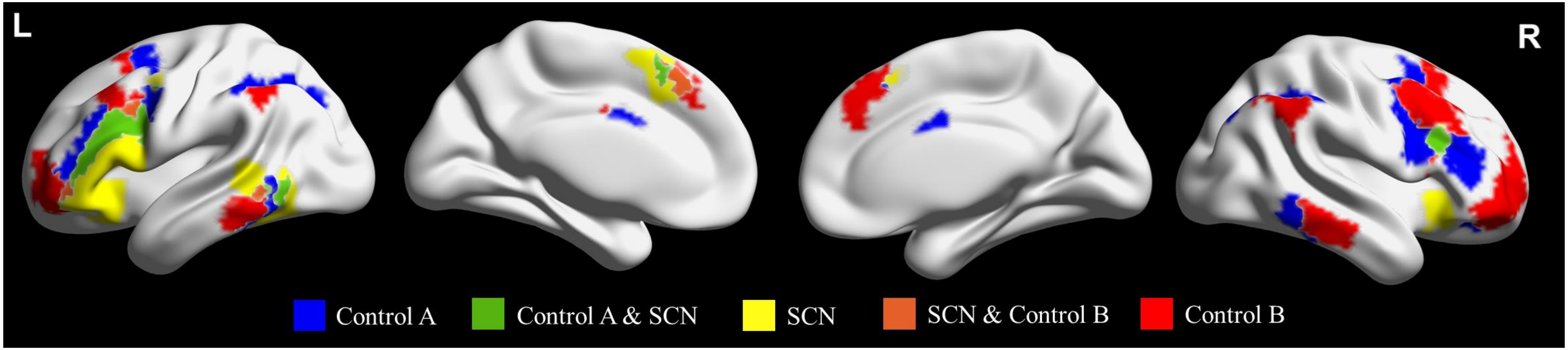


*Supplementary Figure 10. Overlap between SCN, taken from Jackson (2021), and control A and control B, taken from the Yeo et al. (2011) 17-network parcellation. SCN = semantic control network*

The percentage of control A that falls within SCN (16.8%; 1,489/8,848 voxels) is the larger than the percentage of control B that falls within the SCN (5.1%; 508/9,951 voxels). Conversely, a larger percentage of SCN falls within control A (23.4%; 1,489/6,364 voxels) than within control B (8.0%; 508/6,364 voxels).
